# Completion of the gut microbial epi-bile acid pathway

**DOI:** 10.1101/2020.09.27.315549

**Authors:** Heidi L. Doden, Patricia G. Wolf, H. Rex Gaskins, Karthik Anantharaman, João M. P. Alves, Jason M. Ridlon

## Abstract

Bile acids are detergent molecules that solubilize dietary lipids and lipid-soluble vitamins. Humans synthesize bile acids with α-orientation hydroxyl groups which can be biotransformed by gut microbiota to toxic, hydrophobic bile acids, such as deoxycholic acid (DCA). Gut microbiota can also convert hydroxyl groups from the α-orientation through an oxo-intermediate to the β-orientation, resulting in more hydrophilic, less toxic bile acids. This interconversion is catalyzed by regio-(C-3 vs. C-7) and stereospecific (α vs. β) hydroxysteroid dehydrogenases (HSDHs). So far, genes encoding the urso-(7α-HSDH & 7β-HSDH) and iso-(3α-HSDH & 3β-HSDH) bile acid pathways have been described. Recently, multiple human gut clostridia were reported to encode 12α-HSDH, which interconverts DCA and 12-oxolithocholic acid (12-oxoLCA). 12β-HSDH completes the epi-bile acid pathway by converting 12-oxoLCA to the 12β-bile acid denoted epiDCA; however, gene(s) encoding this enzyme have yet to be identified. We confirmed 12β-HSDH activity in cultures of *Clostridium paraputrificum* ATCC 25780. From six candidate *C. paraputrificum* ATCC 25780 oxidoreductase genes, we discovered the first gene (DR024_RS09610) encoding bile acid 12β-HSDH. Phylogenetic analysis revealed unforeseen diversity for 12β-HSDH, leading to validation of two additional bile acid 12β-HSDHs through a synthetic biology approach. By comparison to a previous phylogenetic analysis of 12α-HSDH, we identified the first potential C-12 epimerizing strains: *Collinsella tanakaei* YIT 12063 and *Collinsella stercoris* DSM 13279. A Hidden Markov Model search against human gut metagenomes located putative 12β-HSDH genes in about 30% of subjects within the cohorts analyzed, indicating this gene is relevant in the human gut microbiome.

## Introduction

The human liver produces all 14 enzymes necessary to convert cholesterol into the dihydroxy bile acid chenodeoxycholic acid (3α,7α-dihydroxy-5β-cholan-24-oic acid; CDCA) and the trihydroxy bile acid cholic acid (3α,7α,12α-trihydroxy-5β-cholan-24-oic acid; CA).^1^ These bile acids are conjugated to taurine or glycine in the liver helping to lower the *pKa* and maintain solubility, impermeability to cell membranes, and lower the critical micellar concentration, allowing for efficient emulsification of dietary lipids and lipid-soluble vitamins.^2^ Bile acids are effective detergents owing to the α-orientation of the hydroxyl groups which produce a hydrophilic-face above the plane of the cyclopentanephenanthrene steroid nucleus, and a hydrophobic-face below the plane of the hydrocarbon rings.^1^ Conjugated bile acids emulsify lipids throughout the duodenum, jejunum, and ileum. Once bile acids reach the terminal ileum, high affinity transporters (intestinal bile acid transporter, IBAT) actively transport both conjugated and unconjugated bile acids from the intestinal lumen into ileocytes where they are bound to ileal bile acid binding protein (IBABP) and exported across the basolateral membrane into portal circulation and returned to the liver.^3^ This process of recycling of bile acids is known as the enterohepatic circulation (EHC), responsible for recirculating the ∼2 g bile acid pool 8-10 times daily. While ∼95% efficient, roughly 600-800 mg bile acids escape active transport and enter the large intestine.^2^

Anaerobic bacteria adapted to inhabiting the large intestine have evolved enzymes to modify the structure of host bile acids.^2^ Conjugated bile acids are hydrolyzed, releasing the amino acids, by bile salt hydrolases (BSH) in diverse gut bacteria representing the major phyla, including Bacteroidetes, Firmicutes, and Actinobacteria as well as the domain Archaea.^4^ By contrast, the unconjugated primary bile acids CA and CDCA are 7α-dehydroxylated by a select few species of gram-positive Firmicutes mostly in the genus *Clostridium*, forming deoxycholic acid (3α,12α-dihydroxy-5β-cholan-24-oic acid; DCA) and lithocholic acid (3α-hydroxy-5β-cholan-24-oic acid; LCA), respectively.^1, 5^ The secondary bile acids DCA and LCA have increased hydrophobicity relative to their primary counterparts, which is associated with elevated toxicity.^6^ DCA and LCA have been causally linked to cancers of the colon^7^, liver^8^, and esophagus^9^. Importantly, gut microbiota can produce less toxic oxo-bile acids and β-hydroxy bile acids as well.^6^

Bile acid 3α-, 7α-, and 12α-hydroxyl groups can be reversibly oxidized and epimerized to the β-orientation by pyridine nucleotide-dependent hydroxysteroid dehydrogenases (HSDHs) distributed across the major phyla including Firmicutes, Bacteroidetes, Actinobacteria, Proteobacteria, as well as methanogenic archaea.^1, 10^ HSDH enzymes that recognize bile acids are regio- (C-3 vs. C-7) and stereospecific (α vs. β) for hydroxyl groups decorating the steroid nucleus. Thus, bile acid 12α-HSDH reversibly converts the C-12 position of bile acids from the α-orientation, such as on DCA, to 12-oxo bile acids, such as 12-oxolithocholic acid (12-oxoLCA).^11–14^ Bile acid 12β-HSDH completes the epimerization by interconverting 12-oxo bile acids to the 12β-configuration, forming epi-bile acids. We recently identified and characterized NAD(H)- and NADP(H)-dependent 12α-HSDHs from *Eggerthella* sp. CAG:298^15^, *Clostridium scindens, C. hylemonae*, and *Peptacetobacter hiranonis* (formerly *Clostridium hiranonis*)^10^. In addition to these recently reported 12ɑ-HSDHs, multiple genes encoding enzymes in the urso-(7α-& 7β-HSDH) and iso- (3α- & 3β-HSDH) bile acid pathways have been described to date (**Figure 1**).^5^ However, a gene encoding 12β-HSDH to complete the epi-bile acid pathway has not yet been reported.

**Figure 1.**
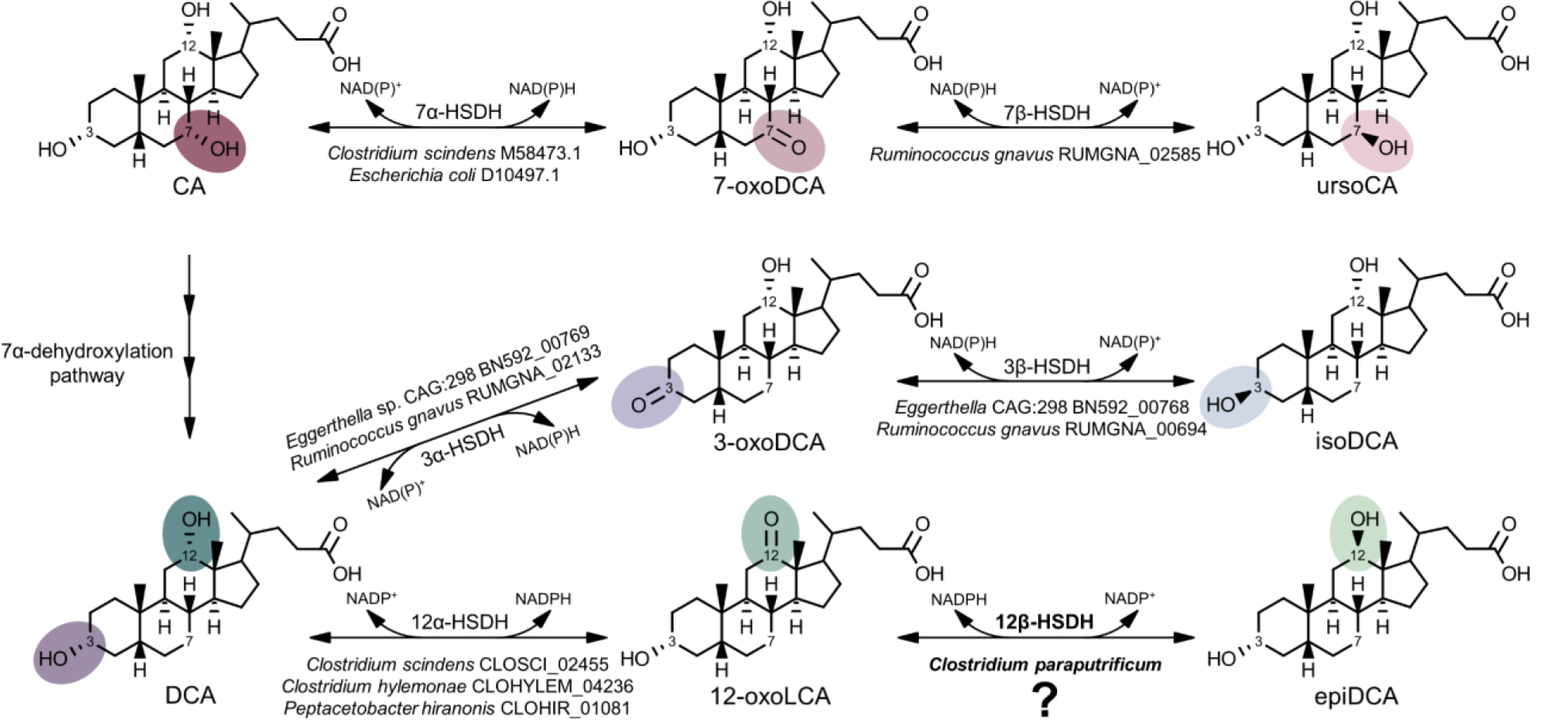
A gene encoding 12β-HSDH completes the gut microbial epi-bile acid pathway. Cholic acid (CA) is converted to the oxo-intermediate, 7-oxodeoxycholic acid (7-oxoDCA), and further to ursoCA through the urso-bile acid pathway catalyzed by NAD(P)-dependent 7α- and 7β- HSDH. The secondary bile acid deoxycholic acid (DCA) is formed through the multi-step 7α- dehydroxylation of CA. DCA is biotransformed to 3-oxoDCA by 3α-HSDH and to isoDCA by 3β-HSDH in the iso-bile acid pathway. DCA can be converted to 12-oxolithocholic acid (12- oxoLCA) by 12α-HSDH and from 12-oxoLCA to epiDCA by 12β-HSDH. Examples of bacteria expressing each HSDH are shown below the reaction followed by corresponding gene annotations. Prior to this study, a gene encoding 12β-HSDH had not been identified.

The first indication that gut bacteria may encode 12β-HSDH was suggested by the detection of 12β-hydroxy bile acids in human feces.^16–18^ Edenharder and Schneider (1985) reported 12β-dehydrogenation of bile acids by *Clostridium paraputrificum*, and epimerization of DCA by co-culture with *E. lenta* and *C. paraputrificum*.^19^ Thereafter, Edenharder and Pfützner (1988) characterized crude NADP(H)-dependent 12β-HSDH activity from *C. paraputrificum* D 762-06.^20^ However, little is known about the potential diversity of gut bacteria capable of forming 12β- hydroxy bile acids that molecular analysis is predicted to yield. Here, we report the identification of a gene encoding NADP(H)-dependent 12β-HSDH from *C. paraputrificum* ATCC 25780 and characterization of the recombinant gene products purified after heterologous expression in *E. coli* from *C. paraputrificum.* We also identify novel taxa encoding bile acid 12β-HSDH by phylogenetic analysis, confirmed by a synthetic biology approach.

## Results

### C. paraputrificum ATCC 25780 possesses bile acid 12β-HSDH activity

We first investigated the bile acid metabolizing capability of *C. paraputrificum* ATCC 25780 because previous studies reported bile acid 12β-HSDH activity in other *C. paraputrificum* strains, but did not identify the gene(s) responsible.^20^ The epi-bile acid pathway of DCA involves the reversible conversion of DCA (3α,12α) to 12-oxoLCA (3α,12-oxo) through the action of 12α-HSDH, and 12-oxoLCA to epiDCA (3α,12β) by 12β-HSDH (**Figure 1**). *C. paraputrificum* ATCC 25780 was incubated with two potential substrates of 12β-HSDH, 12-oxoLCA and epiDCA, along with DCA as a control. In order to contrast the product formed by bile acid 12β-HSDH with that formed by bile acid 12α-HSDH activity, *Clostridium scindens* ATCC 35704, which is known to express 12α-HSDH, was incubated with the same substrates. When 12-oxoLCA was incubated in cultures of *C. paraputrificum* ATCC 25780, the primary product eluted at 13.97 min with 391.28 m/z in negative ion mode (**Figure 2**). This is consistent with the elution time of epiDCA standard and its 392.57 amu formula weight. With epiDCA as substrate, the culture produced a major peak of 391.28 at 13.96 min and a minor peak of 389.27 m/z at 14.34 min, which suggests epiDCA was not converted in large quantities to 12-oxoLCA (390.56 amu). *C. paraputrificum* incubation with DCA did not result in detectable formation of 12-oxoLCA or epiDCA products. Taken together, these data demonstrate *C. paraputrificum* ATCC 25780 expresses bile acid 12β-HSDH activity, but not bile acid 12α-HSDH. *C. scindens* ATCC 35704 incubation with 12-oxoLCA resulted in a main product (15.57 min and 391.28 m/z) consistent with DCA (392.57 amu), demonstrating bile acid 12α-HSDH activity. In addition, reaction with DCA resulted in a peak at 15.57 min and 391.28 m/z along with a peak agreeing with 12-oxoLCA at 14.34 min and 389.27 m/z. When epiDCA was incubated with cultures of *C. scindens* ATCC 35704, we did not observe formation of 12-oxoLCA.

**Figure 2.**
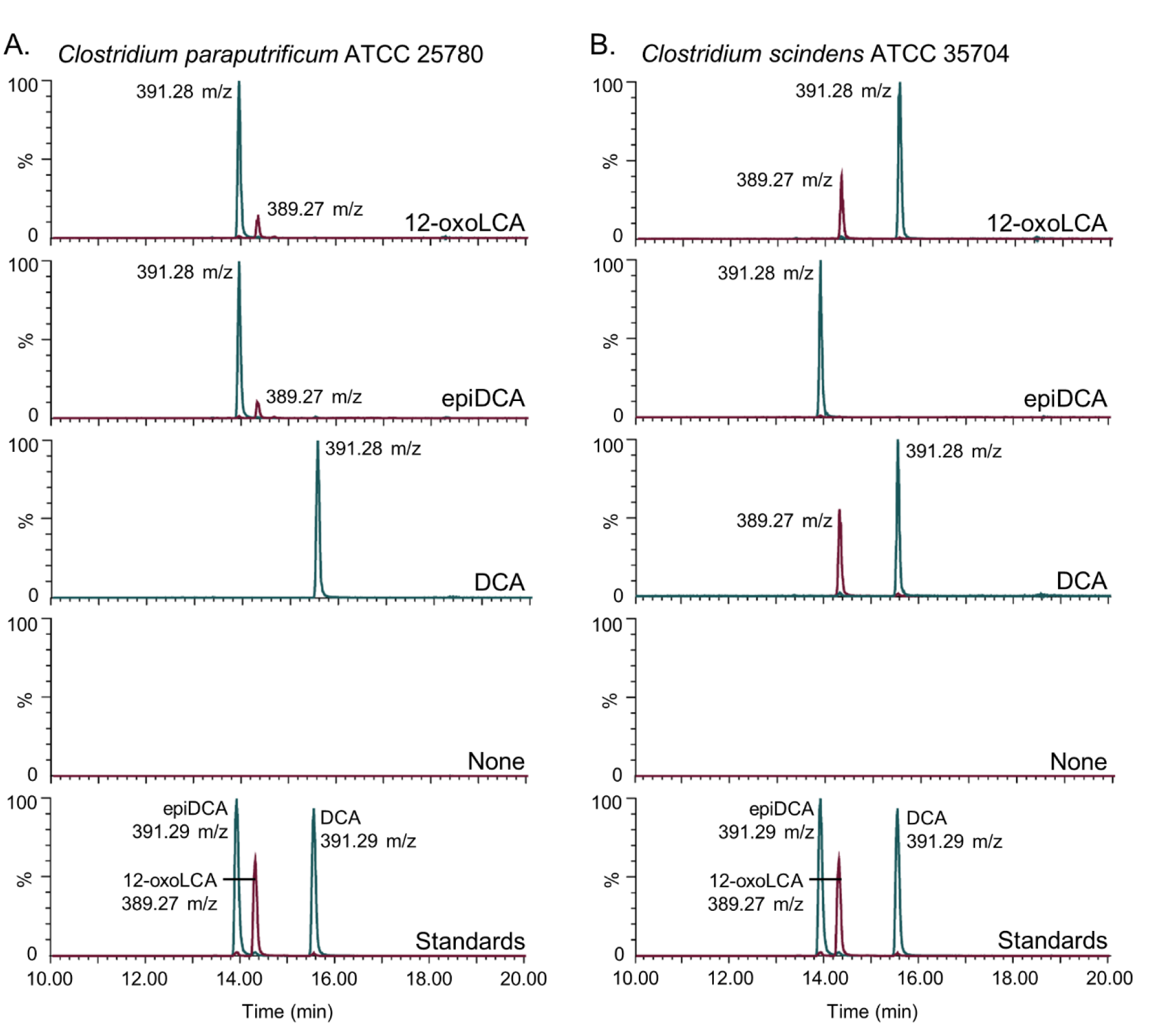
*Clostridium paraputrificum* ATCC 25780 expresses 12β-HSDH while *C. scindens* ATCC 35704 expresses 12α-HSDH by whole-cell LC-MS. (A) Representative negative ion mode LC-MS chromatograms in single ion monitoring mode overlaid with linked vertical axes of *C. paraputrificum* reaction products from 50 μM substrate compared to deoxycholic acid (DCA), 12-oxolithocholic acid (12-oxoLCA) and epiDCA standards. (B) As a control, representative negative ion mode LC-MS chromatograms in single ion monitoring mode overlaid with linked vertical axes of *C. scindens* products from 50 μM substrate was used to demonstrate 12α-HSDH activity. Standards are shown in A and B for ease of comparison to products. Formula weight for DCA is 392.57 atomic mass units (amu), 12-oxoLCA is 390.56 amu, epiDCA is 392.57 amu.

### Identification of a gene encoding bile acid 12β-HSDH

After bile acid 12β-HSDH activity was confirmed in *C. paraputrificum* ATCC 25780, its genome was searched for genes encoding proteins annotated as oxidoreductases within the NCBI database. HSDHs are NAD(P)-dependent and often members of the large and diverse SDR (short-chain dehydrogenase/reductase) family.^21^ Five SDR family oxidoreductase proteins and one aldo/keto reductase were identified as 12β-HSDH candidates in the *C. paraputrificum* ATCC 25780 genome and pursued for further study. These six genes were amplified from genomic DNA of *C. paraputrificum* ATCC 25780, cloned into the pET-28a(+) vector, and overexpressed in *E. coli* (**Table S1**). The N-terminal His6-tagged recombinant proteins were purified by metal-affinity chromatography and resolved by SDS-PAGE (**Figure 3A**).

**Figure 3.**
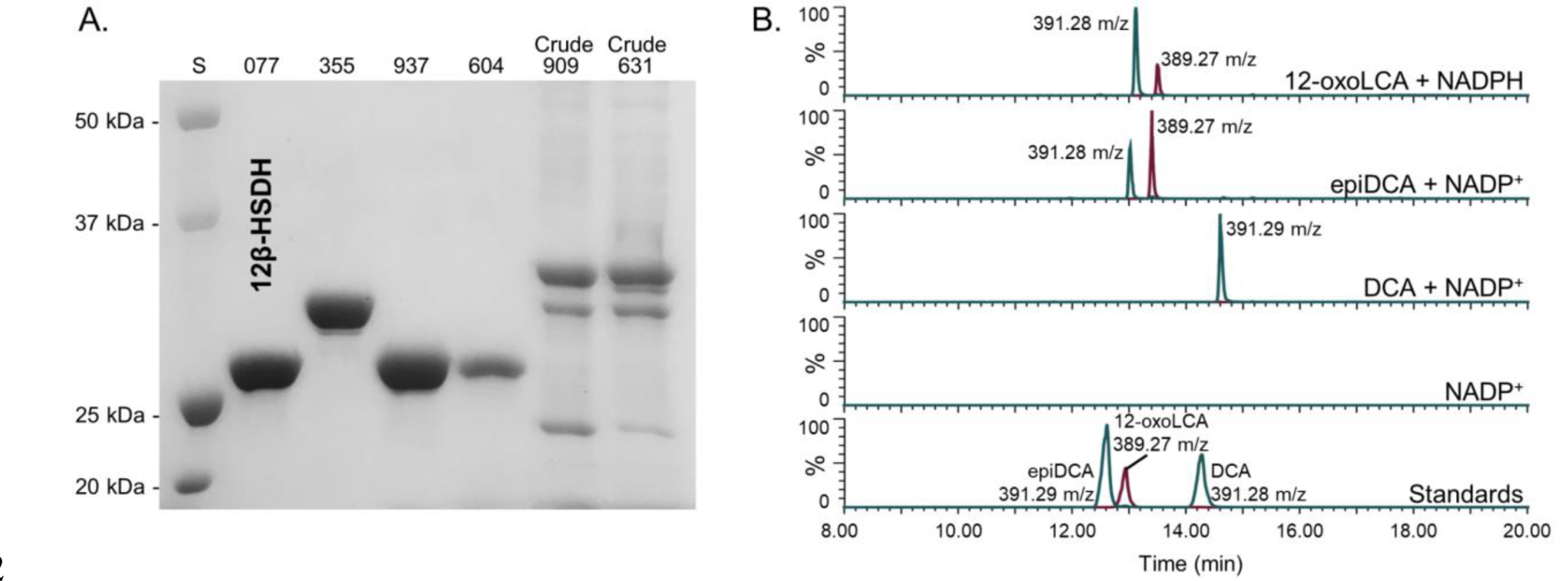
Identification of a gene encoding 12β-HSDH. (A) SDS-PAGE of candidate *Clostridium paraputrificum* 12β-HSDH proteins that were heterologously expressed in *E. coli* and purified with TALON® metal affinity resin. Lanes are as follows: S, molecular weight protein standard; 077, WP_027099077.1; 355, WP_027098355.1; 937, WP_027097937.1; 604, WP_027098604.1; 909, WP_027096909.1; 631, WP_027099631.1. (B) Representative negative ion mode LC-MS chromatograms in single ion monitoring mode overlaid with linked vertical axes of WP_027099077.1 reaction products compared to deoxycholic acid (DCA), 12-oxolithocholic acid (12-oxoLCA) and epiDCA standards. Standards were run on a separate day and show a slight offset in elution time. Reactions consisted of 10 nM WP_027099077.1 with 50 μM (or no) substrate, 150 μM pyridine nucleotide in 50 mM sodium phosphate, 150 mM sodium chloride buffer at pH 7.0. Formula weight for DCA is 392.57 atomic mass units (amu), 12-oxoLCA is 390.56 amu, epiDCA is 392.57 amu.

**Table 1.** Steady-state kinetic parameters of purified recombinant 12β-HSDH.

| Enzyme | Kinetic parameter | Substrate or cofactor<sup>a</sup> |  |  |  |
| --- | --- | --- | --- | --- | --- |
|  |  | 12-oxoLCA<sup>b</sup> | NADPH | epiDCA | NADP<sup>+</sup> |
| Cp12β-HSDH | $K_m$ (μM) | 18.76 ± 0.40<sup>c</sup> | 29.16 ± 0.42 | 44.43 ± 1.13 | 36.84 ± 0.55 |
| | $k_{cat}$ (s<sup>-1</sup>) | 26.62 ± 1.62 | 28.15 ± 1.74 | 59.32 ± 5.47 | 44.61 ± 3.77 |
| | $V_{max}$ (μmol · min<sup>-1</sup> · mg<sup>-1</sup>) | 58.41 ± 3.56 | 61.75 ± 3.82 | 130.13 ± 12.01 | 97.87 ± 8.27 |
| | $k_{cat}/K_m$ (μM<sup>-1</sup> · s<sup>-1</sup>) | 1.42 ± 0.11 | 0.97 ± 0.07 | 1.34 ± 0.15 | 1.21 ± 0.12 |

<sup>a</sup> Assays were performed at saturating concentrations of components not being tested (refer to Materials and Methods).
<sup>b</sup> 12-oxolithocholic acid (12-oxoLCA), epideoxycholic acid (epiDCA).
<sup>c</sup> Values represent the mean ± SD of three or more replicates.

Two out of the six recombinant proteins (WP_027096909.1, WP_027099631.1) were not soluble and bands at the expected molecular masses were apparent in the membrane fraction by SDS-PAGE. These proteins were not explored further. The other four 12β-HSDH candidates (WP_027099077.1, WP_027098355.1, WP_027097937.1, WP_027098604.1) were soluble and visualized by SDS-PAGE. The four soluble recombinant proteins were then screened for pyridine nucleotide-dependent bile acid 12β-HSDH activity by TLC and spectrophotometric assay. Screening reactions were prepared with 12-oxoLCA and NADPH, or epiDCA and NADP^+^ in pH 7.0 phosphate buffer.

Only WP_027099077.1 exhibited 12β-HSDH activity by TLC and spectrophotometric assay, which was also confirmed by LC-MS (**Figure 3B**). Reaction products of WP_027099077.1 with 12-oxoLCA and NADPH, epiDCA and NADP^+^, DCA and NADP^+^, and no substrate control were subjected to LC-MS. In the presence of purified recombinant WP_027099077.1 and NADPH, 12-oxoLCA was reduced quantitatively (2 hydrogen addition) to a product that eluted at 13.12 min with 391.28 m/z in negative ion mode. This is consistent with the 392.57 amu formula weight and elution time for epiDCA based on the substrate standard. Additionally, epiDCA was oxidized to a product with an elution time of 13.40 min at 389.27 m/z, agreeing with the retention time and formula weight of 390.56 amu for authentic 12-oxoLCA. DCA (392.57 amu) was not converted by WP_027099077.1 as the sole peak observed matched DCA standard at 14.60 min and 391.29 m/z. The interconversion of 12-oxoLCA and epiDCA, but no activity with DCA, indicates stereospecificity for the 12β-hydroxy position. Thus, DR024_RS09610 has been identified as the first gene reported that encodes bile acid 12β-HSDH (WP_027099077.1).

Recombinant *C. paraputrificum* WP_027099077.1, hereafter referred to as Cp12β-HSDH, had a theoretical subunit molecular mass of 27.4 kDa. The observed subunit molecular mass was 26.4 ± 0.5 kDa by SDS-PAGE, calculated from three independent protein gels. WP_027099077.1 is predicted to be a cytosolic protein that is not membrane-associated by TMHMM v. 2.0.^22^

### Biochemical characterization of recombinant Cp12β-HSDH

The approximate native molecular mass of Cp12β-HSDH was determined by size-exclusion chromatography. Cp12β-HSDH exhibited an elution volume of 15.04 ± 0.02 mL, corresponding to a 54.67 ± 0.79 kDa molecular mass relative to protein standards (**Figure 4A**). The size-exclusion data along with the theoretical subunit molecular mass of 27.4 kDa suggests Cp12β-HSDH assembles a homodimeric quaternary structure in solution. In order to optimize the enzymatic activity of Cp12β-HSDH, the conversion of pyridine nucleotides at 340 nm was measured in buffers between pH 6.0 and 8.0 by spectrophotometric assay (**Figure 4B**). The optimum pH for Cp12β-HSDH in the oxidative direction with epiDCA as the substrate and NADP^+^ as co-substrate was pH 7.5. In the reductive direction where 12-oxoLCA was the substrate and NADPH the co-substrate, the optimum pH was 7.0.

**Figure 4.**
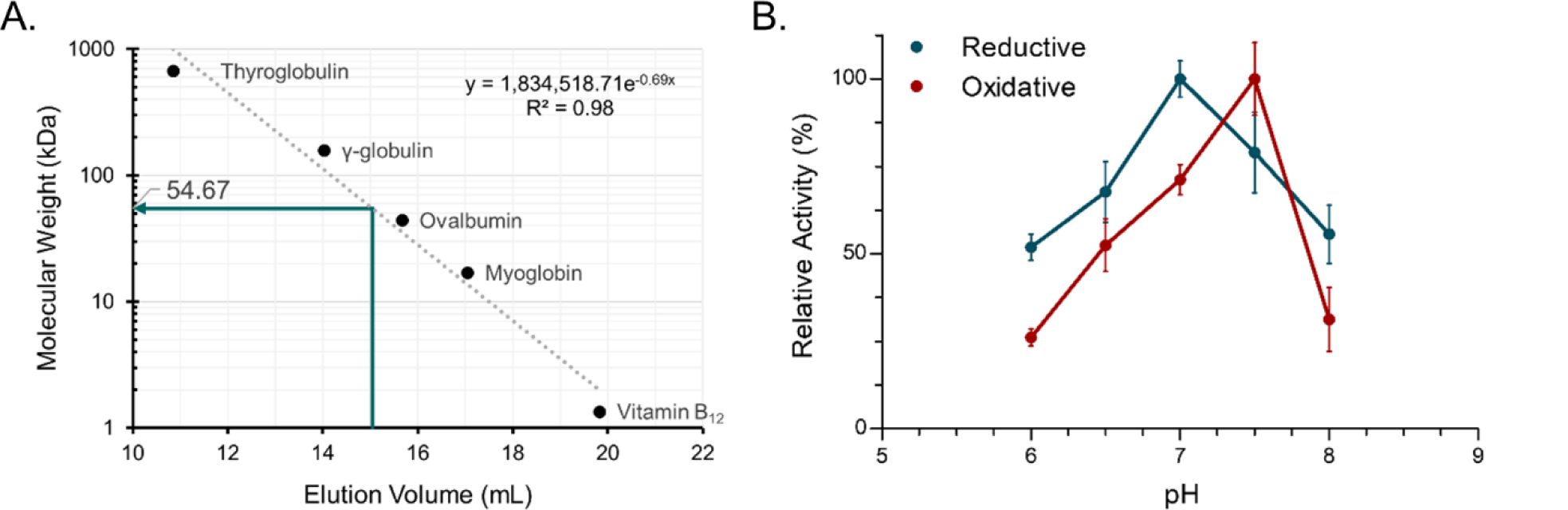
Biochemical characterization of recombinant *C. paraputrificum* 12β-HSDH. (A) Native molecular size analysis of 10 mg/mL purified 12β-HSDH via size-exclusion chromatography. (B) Effect of pH on 12β-HSDH activity. The reaction in the reductive direction (blue) consisted of 12-oxoLCA as substrate with NADPH as cofactor. The oxidative reaction (red) was epiDCA with NADP^+^. See Materials and Methods for buffer compositions.

Michaelis-Menten kinetics were performed at the pH optimum for each direction. In the reductive direction, Cp12β-HSDH displayed a *Km* value for 12-oxoLCA at 18.76 ± 0.40 µM which was similar to that of NADPH (**Table 1**; **Figure S1**). The *Km* value in the oxidative direction with epiDCA as substrate was about twice the *Km* determined for 12-oxoLCA. The *Km* for NADP^+^ was 36.84 ± 0.55 µM. The *Vmax* and *kcat* were greater in the oxidative than the reductive direction.

However, the catalytic efficiency (*kcat/Km*) of 12-oxoLCA as substrate was greater than the oxidative direction with epiDCA as substrate.

Pyridine nucleotide cofactor and bile acid substrate-specificity of Cp12β-HSDH were determined by relative activity compared to either 12-oxoLCA or epiDCA through spectrophotometric assay (**Table 2**). NAD^+^ and NADH were not co-substrates for Cp12β-HSDH. DCA (3α,12α) as well as CA (3α,7α,12α) were not substrates, which is expected because they are 12α-hydroxy bile acids not 12β-hydroxy bile acids. CDCA (chenodeoxycholic acid; 3α,7α) lacks a 12-hydroxyl group, and as expected was not a substrate. The CA derivatives 12-oxoCDCA (3α,7α,12-oxo) and epiCA (3α,7α,12β) had ∼12% and 27% activity, respectively, relative to bile acids lacking a 7α-hydroxyl group. The activity of 3,12-dioxoLCA was ∼19% compared to 12-oxoLCA. Altogether, the results suggest Cp12β-HSDH is specific for NADP(H) and favors 12-oxoLCA and epiDCA over their 7α-hydroxy counterparts.

**Table 2.** Substrate and pyridine nucleotide specificity of purified recombinant *C. paraputrificum* 12β-HSDH, *Eisenbergiella* WP_118677302.1, *Olsenella* WP_120179297.1, and *Novosphingobium* WP_007678535.1.

| Enzyme | Substrate<sup>a,b</sup> | Cofactor | Activity (μmol·min<sup>-1</sup>·mg<sup>-1</sup>) | Relative activity (%) |
| --- | --- | --- | --- | --- |
| Cp12β-HSDH | 12-oxoLCA | NADPH | 18.26 ± 1.01<sup>c</sup> | 100 |
|  | 12-oxoLCA | NADH | -<sup>d</sup> | - |
| | 12-oxoCDCA | NADPH | 2.21 ± 0.82 | 12.08 |
| | 3,12-dioxoLCA | NADPH | 3.49 ± 0.34 | 19.09 |
| WP_118677302.1 | 12-oxoLCA | NADPH | 16.04 ± 1.23 | 87.85 |
| WP_120179297.1 | 12-oxoLCA | NADPH | 23.29 ± 2.57 | 127.57 |
| WP_007678535.1 | 12-oxoLCA | NADPH | - | - |
| Cp12β-HSDH | epiDCA | NADP<sup>+</sup> | 33.42 ± 0.81 | 100 |
|  | epiDCA | NAD<sup>+</sup> | - | - |
| | epiCA | NADP<sup>+</sup> | 8.99 ± 0.90 | 26.88 |
|  | DCA | NADP<sup>+</sup> | - | - |
|  | CA | NADP<sup>+</sup> | - | - |
|  | CDCA | NADP<sup>+</sup> | - | - |
| WP_118677302.1 | epiDCA | NADP<sup>+</sup> | 27.85 ± 1.12 | 83.32 |
| WP_120179297.1 | epiDCA | NADP<sup>+</sup> | 23.02 ± 2.57 | 68.86 |
| WP_007678535.1 | epiDCA | NADP<sup>+</sup> | - | - |

<sup>a</sup> 12-oxolithocholic acid (12-oxoLCA), 12-oxochenodeoxycholic acid (12-oxoCDCA), deoxycholic acid (DCA), cholic acid (CA).
<sup>b</sup> Assays were performed with 10 nM enzyme, 50 μM substrate and 150 μM cofactor at optimum pH.
<sup>c</sup> Values represent the mean ± SD of three or more replicates.
<sup>d</sup> -, no activity detected.

### Phylogenetic analysis of Cp12β-HSDH

The Cp12β-HSDH sequence from *C. paraputrificum* ATCC 25780 (WP_027099077.1) was used in a BLASTP search against the NCBI non-redundant protein database in order to determine its prevalence across bacteria. A maximum likelihood phylogeny of 5,000 sequences was constructed, revealing that many sequences most similar to Cp12β-HSDH are found in Firmicutes and Actinobacteria (**Figure S2**). Within the 5,000-member phylogeny, a subtree (highlighted grey) of the most closely related proteins to Cp12β-HSDH was selected for closer inspection (**Figure 5**). Cp12β-HSDH clustered most closely with other *C. paraputrificum* sequences (WP_099327725, WP_049179624, WP_111937163). These sequences are encoded by *C. paraputrificum s*trains isolated from preterm infants, namely strain LH025, LH141, and LH058^23^, or isolated from feces (Gcol.A11).^24^

**Figure 5.**
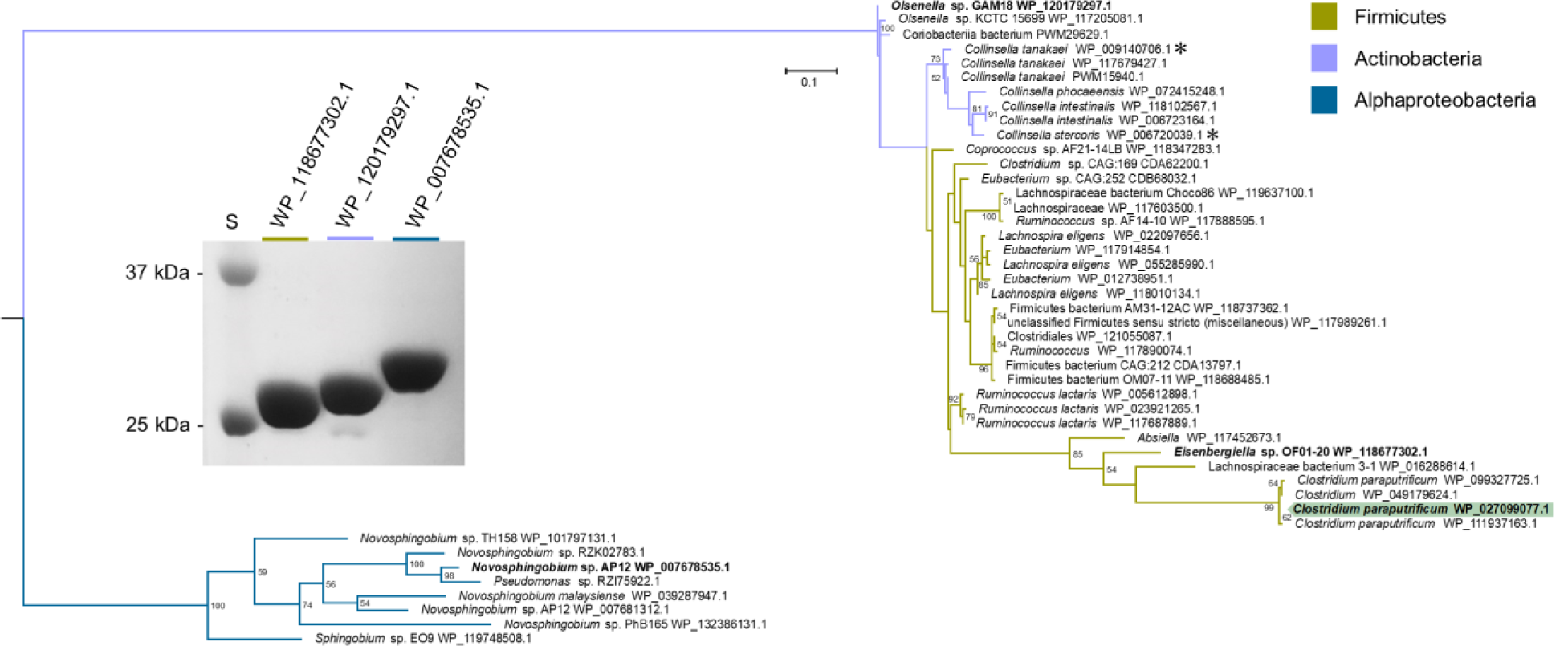
Maximum-likelihood tree based on a subset of the taxa present in the full phylogenetic analysis of 12β-HSDH and SDS-PAGE of proteins explored further. Sequences selected for this analysis were those nearest to the *C. paraputrificum* 12β-HSDH (highlighted), plus an outgroup. For the full tree with about 5,000 sequences, see Figure S1. Taxonomic affiliations are indicated by branch colors as specified in the legend. Bolded sequences were chosen for further study. Asterisks indicate novel C-12 epimerizing organisms. (Inset) SDS-PAGE of purified recombinant *Eisenbergiella* WP_118677302.1, *Olsenella* WP_120179297.1, and *Novosphingobium* WP_007678535.1 heterologously expressed in *E. coli* and purified with TALON® metal affinity resin. S, molecular weight protein standard.

Firmicutes harbor the majority of sequences within the 12β-HSDH subtree, spanning genera including *Eisenbergiella*, *Ruminococcus*, and *Coprococcus*. To determine if other organisms within the tree have bile acid 12β-HSDH activity, the gene encoding WP_118677302.1 from *Eisenbergiella* sp. OF01-20 was synthesized by IDT in the codon-usage of *E. coli* (**Table S1**), cloned into pET-28a(+), overexpressed in *E. coli*, and purified by affinity chromatography (**Figure 5**). Recombinant WP_118677302.1 was screened by spectrophotometric assay with NAD(P)(H) against 12-oxoLCA, epiDCA, and DCA. Relative to Cp12β-HSDH, WP_118677302.1 exhibited 88% activity with 12-oxoLCA, and 83% activity with epiDCA. WP_118677302.1 did not show conversion of DCA, confirming that this enzyme has bile acid 12β-HSDH activity (**Table 2**).

The subtree also contains many sequences from Actinobacteria, the genera *Collinsella* and *Olsenella* among them. C*ollinsella* species are of interest because *C. aerofaciens* expresses BSH and various HSDHs recognizing sterols^25^, including bile acid 12α-HSDH.^26^ To determine if a member of the Actinobacteria encodes a bile acid 12β-HSDH, a sequence more distantly related to Cp12β-HSDH, *Olsenella* sp. GAM18 WP_120179297.1, was chosen for gene synthesis and protein overexpression because it had not been shown previously to metabolize bile acids (**Figure 5**). 12-oxoLCA, epiDCA and DCA were tested as substrates and conversion was measured by spectrophotometric assay. Recombinant WP_120179297.1 displayed activity with 12-oxoLCA at 128% relative to Cp12β-HSDH, 69% relative activity with epiDCA, and showed no reaction with DCA (**Table 2**). These data confirm that the more distantly related WP_120179297.1 has bile acid 12β-HSDH activity.

Within the extended subtree are various *Novosphingobium s*pecies. These Alphaproteobacteria deserve mention due to their ability to biodegrade aromatic compounds, such as phenanthrene^27^ and estrogen.^28^ To test if this cluster has bile acid 12β-HSDH activity, WP_007678535.1 from *Novosphingobium* sp. AP12 was synthesized, cloned, overexpressed, and purified (**Figure 5**). The potential 12β-HSDH activity of WP_007678535.1 was screened using 12-oxoLCA, epiDCA, and DCA as substrates. WP_007678535.1 exhibited no activity with these bile acid substrates (**Table 2**). Because *Novosphingobium s*trains are frequently plant-associated or isolated from aquatic environments^29^, this enzyme may be specific for other substrates.

The genomic context of 12β-HSDH genes from *C. paraputrificum* ATCC 25780, *Eisenbergiella* sp. OF01-20, and *Olsenella* sp. GAM18 was explored (**Figure S3**). The three 12β-HSDH genes did not appear to be organized within an operon nor was the genomic context conserved across these organisms.

Two organisms present in the 12β-HSDH subtree, *Collinsella tanakaei* and *Collinsella stercoris* (**Figure 5**, asterisks), were also found in a previous phylogenetic analysis of putative 12α-HSDHs (Doden AEM 2018). Due to strain variation within species, we inspected the sequences further and determined that the pairs of HSDHs are encoded by the same strain within each species. *Collinsella tanakaei* YIT 12063 12α-HSDH (WP_009141301.1) and 12β-HSDH (WP_009140706.1) are encoded by the genes HMPREF9452_RS06335 and HMPREF9452_RS03390, respectively. *Collinsella stercoris* DSM 13279 also contains both putative 12α-HSDH (WP_040360544.1; COLSTE_RS02900) and 12β-HSDH (WP_006720039.1; COLSTE_RS01465).^10^ While the paired activity of 12α/12β-HSDH has not been tested in culture, these organisms may be novel epi-epimerizing strains that convert bile acid 12α-hydroxyl groups to the epi-configuration. To our knowledge, these are the first strains identified with C-12 epimerizing ability.

### Hidden Markov Model search of putative 12β-HSDH genes in human gut metagenomes

To understand the distribution of potential 12β-HSDH genes in the human colonic microbiome, a Hidden Markov Model (HMM) search was performed against metagenome assembled genomes (MAGs) from four publicly available cohorts^30–33^ using reference sequences from the 12β-HSDHs characterized in this paper (**Figure 5**). Putative 12β-HSDH genes inferred by HMM search were found in ∼30% of the subjects (198/666) (**Figure 6A**). Twenty-two subjects exhibited two different organisms containing the gene. This gene was found in healthy subjects as well as subjects with the following disease states: colorectal cancer, colorectal adenoma, fatty liver, hypertension, and type 2 diabetes.

**Figure 6.**
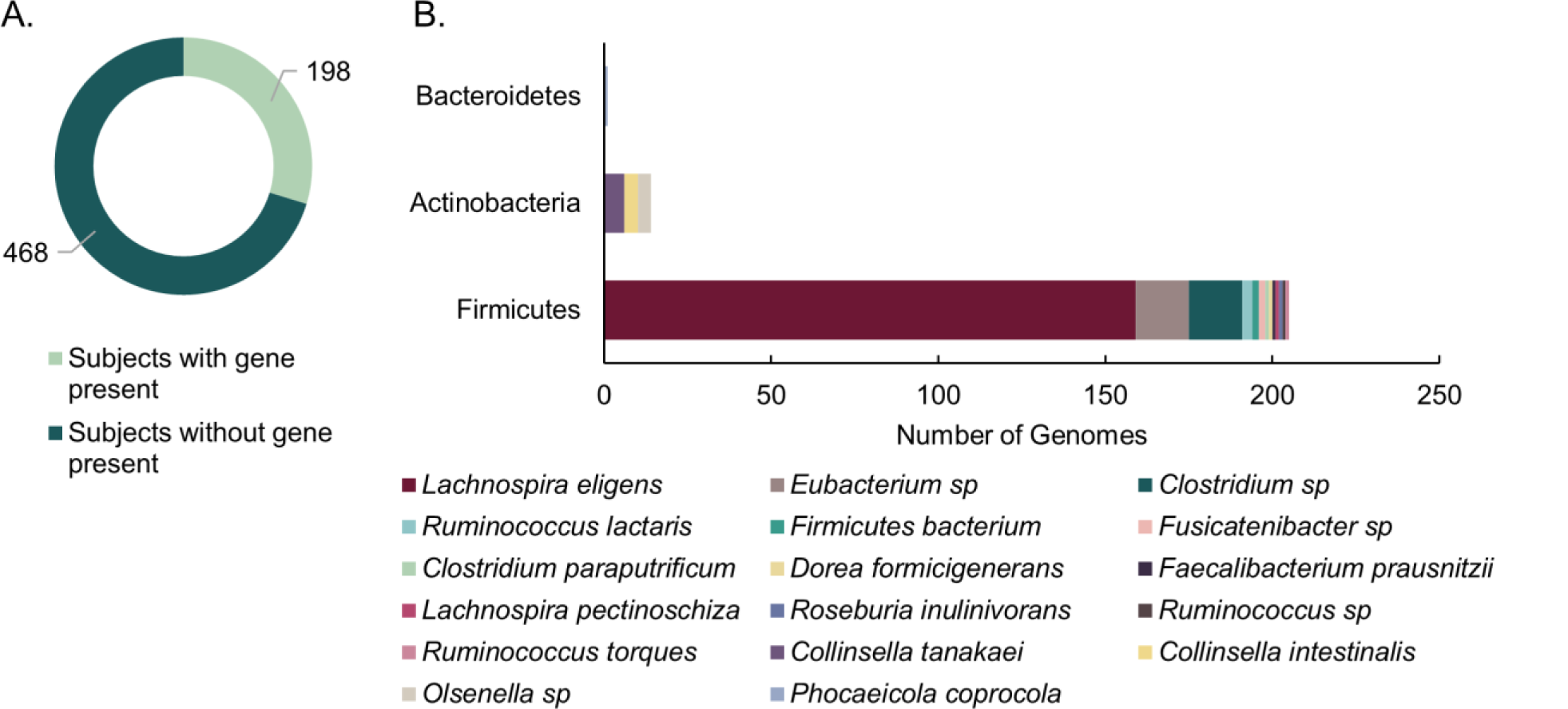
12β-HSDH Hidden-Markov Model search. (A) Number of subjects identified with putative 12β-HSDH genes present in their gut metagenomes. The metagenomes analyzed were from 4 distinct cohorts. (B) Distribution of microbial genomes with putative 12β-HSDH genes present across the 4 metagenomic studies.

Two hundred twenty microbial genomes contained putative 12β-HSDH genes among 16,936 total available genomes. Putative 12β-HSDH genes were most often identified in the phylum Firmicutes, which was dominated by genes in *Lachnospira eligens* (formerly *Eubacterium eligens*) (**Figure 6B**). The gene from *L. eligens* was widespread across subjects in each of the four cohorts. This large proportion of hits from *L. eligens* may reflect its higher relative abundance allowing it to be assembled better into genomes. Sequences from this organism also appeared multiple times in the 12β-HSDH subtree (**Figure 5**). *Lachnospira eligens* is a pectin degrader capable of promoting anti-inflammatory cytokine IL-10 production *in vitro*^34^ and has been proposed as a probiotic for atherosclerosis^35^. The gene was also present in *C. paraputrificum* along with other unidentified *Clostridium* sp. and *Eubacterium* sp. Actinobacteria had few members with the gene, represented by *Collinsella intestinalis*, *Collinsella tanakaei*, and *Olsenella* sp. *Phocaeicola coprocola* (formerly *Bacteroides coprocola*) was the only member of phylum Bacteroidetes with the gene.

### Phylogenetic analysis of regio- and stereospecific HSDHs

Next, the phylogenetic relationship between Cp12β-HSDH (WP_027099077.1) and other regio- and stereospecific HSDHs was explored. To accomplish this, we updated the HSDH phylogeny presented by Mythen et al. (2018) by including additional bacterial or archaeal HSDH sequences of known or putative function along with representative eukaryotic sequences (**Figure 7**; **Table S2**).^15^ The sequences included span the known HSDH functional capacities, with some recognizing bile acids and others recognizing steroids like cortisol or progesterone. Most members of each HSDH class are clustered together, which is apparent by each highlight color encompassing more than one HSDH of the same known function. Furthermore, most bacterial HSDHs grouped separately from their eukaryotic counterparts.

**Figure 7.**
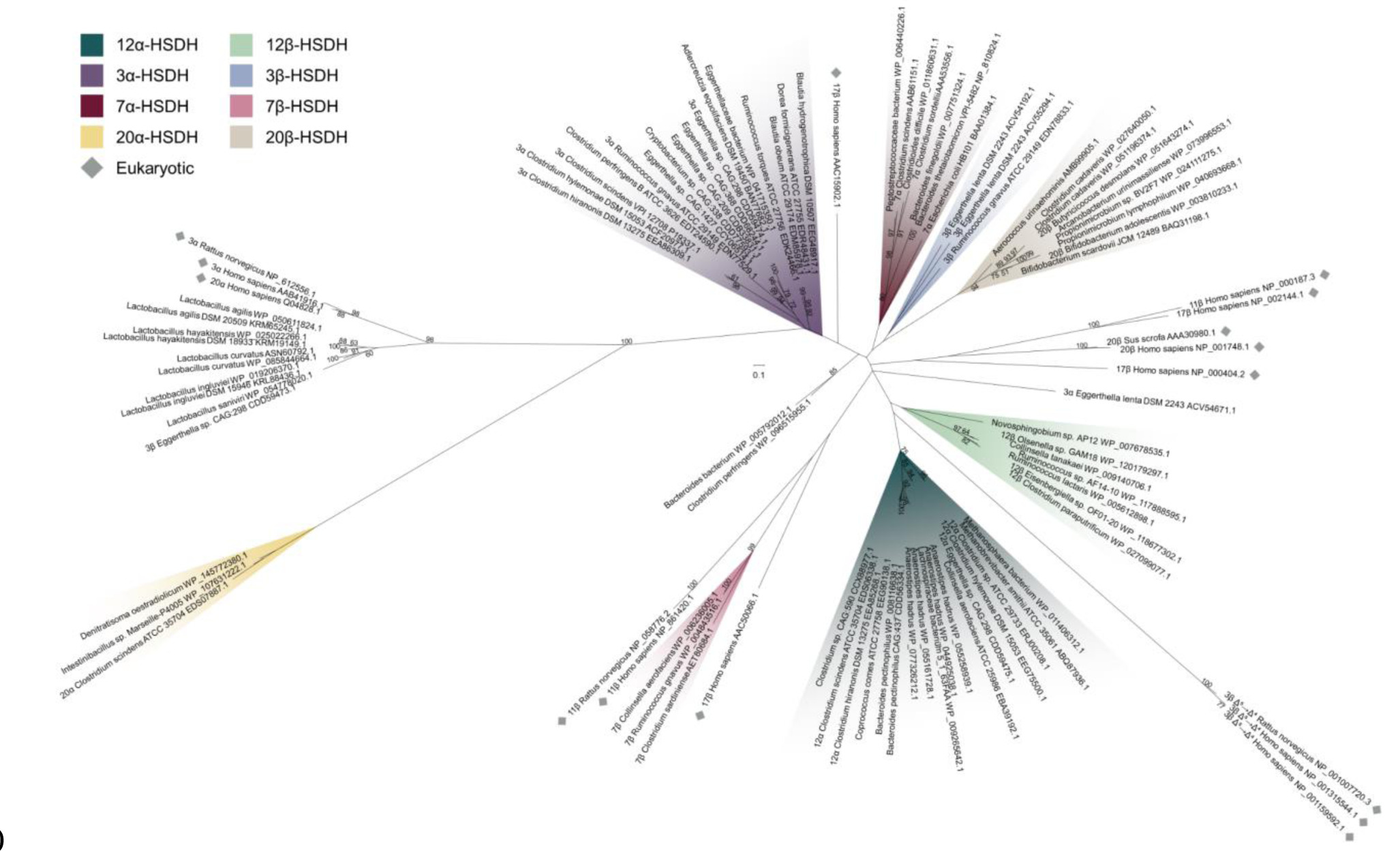
Maximum-likelihood phylogenetic analysis of regio- and stereospecific HSDHs. Clusters are shaded by function or marked as eukaryotic, as displayed in the legend. Sequences with experimentally determined activities are labeled with their function followed by organism and accession number. See Table S2 for sequence information.

Prokaryotic sequences were interspersed among the eukaryotic with some exceptions in grouping by HSDH function. Cp12β-HSDH, the two other confirmed bile acid 12β-HSDHs (WP-118677302.1, WP_120179297.1), and additional similar sequences from across our bile acid 12β-HSDH subtree formed their own cluster. These sequences shared a branch with bacterial bile acid 12α-HSDHs as well as eukaryotic 3β-HSD/Δ^5^→Δ^4^/isomerases. Bile acid 12α-HSDH sequences included various clostridia (EDS06338.1, EEG75500.1, EEA85268.1, ERJ00208.1)^10, 36^ and *Eggerthella* (CDD59475.1).^15^ *Collinsella aerofaciens* (EBA39192.1), which has been reported to express bile acid 12α-HSDH activity^26^, grouped with the known bile acid 12α-HSDHs along with two human gut archaeal sequences from *Methanosphaera stadtmanae* and *Methanobrevibacter smithii*.

Clostridial (gram-positive) bile acid 7α-HSDHs (AAB61151.1 etc.)^37^ clustered separately from those expressed by *E. coli* (BAA01384.1)^38^ and those predicted in *Bacteroides* (gram-negative), similarly to the Mythen et. al. (2018) phylogeny. Bile acid 7β-HSDHs did not closely cluster with other classes of bacterial HSDHs. Instead, the nearest neighbours to the three known bile acid 7β-HSDHs (WP_006236005.1, WP_004843516.1, AET80684.1)^25, 39, 40^ included in this tree were eukaryotic steroid 11β- and 17β-HSDs.

Bacterial 3α-HSDHs clustered together, excluding one outlier from *Eggerthella lenta* (ACV54671.1).^41^ Within the bile acid 3α-HSDH group, four enzymes predicted with this function formed a separate branch apart from the confirmed bile acid 3α-HSDHs. Likewise, three known bile acid 3β-HSDHs grouped together (ACV55294.1, ACV54192.1, EDN78833.1)^41^, while one *Eggerthella* sequence was most closely related to putative bile acid 3β-HSDHs from *Lactobacillus* spp. identified by BLAST search in Mythen et. al (2018).

Bacterial steroid 20β-HSDHs convert the glucocorticoid cortisol to 20β-dihydrocortisol. Two experimentally confirmed 20β-HSDHs (WP_003810233.1, WP_051643274.1)^42, 43^ grouped with putative sequences from both gut and urinary tract isolates. To date, only one steroid 20α-HSDH sequence, which interconverts cortisol and 20α-dihydrocortisol, has been reported (*C. scindens* EDS07887.1).^44, 45^ Therefore, we performed a BLASTP search and found two sequences with high similarity, WP_145772308.1 from *Denitratisoma oestradiolicum* DSM 16959 and WP_107631222.1 from *Intestinibacillus* sp. Marseille-P4005. *D. oestradiolicum* DSM 16959 was isolated from sludge in a municipal wastewater treatment plant and can use 17β-estradiol as a sole carbon and energy source.^46^ *Intestinibacillus massiliensis* strain Marseille-P3216, a close relative to *Intestinibacillus* sp. Marseille-P4005 found in our tree, was isolated from the human colon and is most closely related to the species *Butyricicoccus desmolans* (formerly *Eubacterium desmolans*) by 16S rRNA gene sequence.^47^ Interestingly, *B. desmolans* ATCC 43058 encodes a 20β-HSDH (WP_051643274.1).^43^

Eukaryotic HSDH sequences, typically denoted HSD, were spread throughout the phylogeny, but generally grouped with like sequences. The 17β- and 11β-HSD sequences did not form a group, instead clustering by type. For example, *Homo sapiens* 11β-HSD type 1 (NP_861420.1) was closely related to *Rattus norvegicus* 11β-HSD type 1 (NP_058776.2) and more distantly related to *Homo sapiens* 11β-HSD type 2 (NP_000187.3). 11β-HSD type 1 and type 2 both interconvert steroids between active and inactive forms, such as cortisol and cortisone.^48^

However, 11β-HSD type 1 primarily acts as a reductase in many tissues while 11β-HSD type 2 functions as a dehydrogenase.^48^

## Discussion

Microbial bile acid HSDHs have been studied since the early 1970s, with much of the original work focusing on 3α- and 7α-HSDHs.^14, 49^ Thereafter, 3β-, 7β- and 12α-HSDH activity was observed in cultures of various microbiota^1^, including *Eggerthella lenta* (formerly *Eubacterium lentum*) which is capable of oxidizing CA and DCA at C-12 and epimerizing bile acids at C-3.^50^ In the mid-1980s, *C. paraputrificum*, *C. tertium*, and *Clostridioides difficile* each in binary cultures with *E. lenta* were shown to epimerize DCA via a 12-oxo-intermediate to epiDCA.^19^ Since then, HSDH genes encoding the iso- and urso-bile acid pathways and 12α-HSDH were identified, but not 12β-HSDH.^5^ In this work, we identified the first bile acid 12β-HSDH gene, completing the microbial epi-bile acid pathway.

Edenharder & Pfützner (1988) initially characterized NADP(H)-dependent 12β-HSDH from crude extracts of the fecal isolate *C. paraputrificum* strain D 762-06, with differing results from our findings.^20^ Gel filtration analysis of crude extract from *C. paraputrificum* strain D 762-06 suggested a molecular mass of 126 kDa; whereas our current work with Cp12β-HSDH from ATCC 25780 is estimated at 54.6 KDa by gel filtration chromatography. The strain used in this study, *C. paraputrificum* ATCC 25780, was also isolated from feces.^51^ It is possible that these are the same NADP(H)-dependent enzymes by sequence from two different strains of *C. paraputrificum* and that the recombinant protein quaternary structure is unstable, resulting in a dimeric form in our study. Alternatively, these bacterial strains may have distinct versions of 12β-HSDH with different amino acid sequences, as we have shown previously for 12α-HSDH from

*Eggerthella lenta*.^15, 52^ Indeed, the 12β-HSDH from *C. paraputrificum* strain D 762-06 was reported to be partially membrane associated, whereas hydropathy prediction by TMHMM v. 2.0 found no evidence of transmembrane domains in Cp12β-HSDH. In addition, pH optima for the conversion of 12-oxoLCA between Cp12β-HSDH (7.0) and the native 12β-HSDH (10.0) from strain D 762-06 differed. Oxidation of epiDCA was optimal at pH 7.5 for Cp12β-HSDH, and reported as pH 7.8 for the crude native enzyme from strain D 762-06.^20^ Further work will be needed to determine if distinct bile acid 12β-HSDHs are present in *C. paraputrificum* strains.

Cp12β-HSDH exhibited a dimeric quaternary structure by size-exclusion chromatography under our experimental conditions. Although future crystallization of Cp12β-HSDH would better illustrate its true polymeric state, HSDHs are often either tetrameric^42, 53^ or dimeric.^54, 55^ Cp12β- HSDH was more specific for bile acids lacking a position 7-hydroxyl group: epiDCA and 12-oxoLCA, over epiCA and 12-oxoCDCA. Cp12β-HSDH also had lower activity with 3,12-dioxoLCA versus 12-oxoLCA. This indicates that both the 7-hydroxyl and 3-oxo groups hinder the ability of Cp12β-HSDH to convert the substrate. An x-ray crystal structure of Cp12β-HSDH may shed light on why this apparent steric hindrance occurs.

Phylogenetic analysis of Cp12β-HSDH coupled with synthetic biological “sampling” and validation at different points along the branches revealed shared 12β-HSDH function among *Eisenbergiella* sp. OF01-20 and *Olsenella* sp. GAM18, lending functional credibility to sequences throughout the subtree (**Figure 5**; **Table 2**). *Eisenbergiella* sp. OF01-20 was originally sequenced from a human gut microbiota cultivation project (Integrated Microbial Genomes [IMG] Genome ID: 2840324701). *Eisenbergiella* spp. are often present at relative abundances of less than 0.1% in human fecal samples.^56, 57^ *Olsenella* sp. GAM18 was initially isolated from humans (IMG Genome ID: 2841219092). The relative abundance of *Olsenella* was shown to be about 2% within the gut microbiome of some individuals.^58^ Our subtree includes more abundant gut taxa such as *Ruminococcus* (relative abundance ∼5%)^59, 60^ and *Collinsella* (relative abundance ∼8%)^59^, as well. Due to limitations in 16S rDNA sequencing depth, it is difficult to conclude if the species in our subtree are found at relevant levels in the human gut or if 12β-HSDH genes are present. Therefore, we performed a HMM search to assess the relative prevalence of 12β-HSDH genes. About 30% of subjects had putative 12β-HSDH genes, indicating the relevance of this gene in the human gut microbiome. The HMM search revealed that 220 microbial genomes out of 16,936 total contained putative 12β-HSDH genes. While concrete prevalence is difficult to establish, putative 12β-HSDH genes are less widespread than the ubiquitous bile-acid metabolizing gene, bile salt hydrolase^4^, which was present in 2,456/16,936 total genomes in these cohorts. These data expand the limited metagenomic work that has focused on bile acid HSDH genes in the human gut.^61^

Two organisms from our 12β-HSDH subtree were also identified in a previous 12α-HSDH phylogeny from Doden et al. (2018). Putative proteins WP_009140706.1 (12β-HSDH) and WP_009141301.1 (12α-HSDH) are both present in *Collinsella tanakaei* YIT 12063 and are encoded by the genes HMPREF9452_RS03390 and HMPREF9452_RS06335, respectively. Similarly, *Collinsella stercoris* DSM 13279 encodes both putative 12β-HSDH (WP_006720039.1; COLSTE_RS01465) and 12α-HSDH (WP_040360544.1; COLSTE_RS02900).^10^ Although the dual 12α/12β-HSDH activity is untested in culture, we predict these strains are novel C-12 epimerizers. Epimerizing strains have been identified for the C-3^19, 41^ and C-7 hydroxyl^1, 62^ positions, however, this is the first indication of bacteria capable of C-12 epimerization.

The sequence WP_007678535.1 from *Novosphingobium* sp. AP12, whose recombinant enzyme product did not exhibit bile acid 12β-HSDH activity with the substrates tested, may be specific for aerobic bile acid degradation products. Environmental microorganisms, such as *Comamonas testosteroni* TA441 and *Pseudomonas* sp. strain Chol1, encode a CA degradation pathway involving conversion of a 12-oxo-intermediate to 7α,12β-dihydroxy-androsta-1,4-diene-3,17-dione (12β-DHADD).^63, 64^ Thus, sequences in the extension of the subtree may have 12β-HSDH activity, but with specificity for side-chain cleaved steroids rather than bile acids.

Indeed, this function joins the vast repertoire of HSDHs already studied in many Firmicutes and Actinobacteria.^1^ Bile acid 12α-HSDH activity has been detected in *Eggerthella* species^15, 52^ in the phylum Actinobacteria and various clostridia^10–12, 36^ in the phylum Firmicutes. Similarly, 3α- and 3β-HSDH are widespread among Firmicutes^1,65^, and 3α-HSDH has also been reported in *Eggerthella* species.^13, 15, 41^ 7α- and 7β-HSDH were shown in numerous Firmicutes^14, 37, 65^ and the Actinobacteria *Collinsella aerofaciens*.^25^ Along with these bile acid-specific HSDHs, the glucocorticoid 20α- and 20β-HSDHs are evident in both Firmicutes^43, 45^ and Actinobacteria such as *Bifidobacterium adolescentis*^66^. Until this study, there were no reports of genes encoding 12β-HSDH and the activity had only been shown in *C. paraputrificum*, *C. tertium* and *C. difficile.*^19^ Thus, our phylogenetic analysis revealed hitherto unknown diversity for bile acid 12β-HSDHs within the Firmicutes and Actinobacteria. Bacteroidetes sequences were notably absent within our 12β-HSDH subtree and only one sequence was identified in our HMM search, although Bacteroidetes have been shown to encode multiple other HSDHs.^1, 49^ Interestingly, *C. tertium* and *C. difficile* enzymes were also not present in our phylogenetic analysis even though this activity has been reported for strains of these clostridia^19^, indicating that genes encoding other forms of bile acid 12β-HSDH are present in the gut microbiome.

The distribution pattern of microbial HSDHs is becoming increasingly clear (**Figures 5 & 7**), although in many cases the evolutionary pressures on gut microbes for encoding particular regio- and stereospecific HSDH enzymes is not clear. As observed with BSH enzymes, the functional importance of HSDHs may be strain-dependent. In some strains, the mere ability to acquire or dispose of reducing equivalents may be important, and the class of enzyme unimportant. Bile acid hydroxylation patterns affect the binding and activation/inhibition of host nuclear receptors.^67^ HSDH enzymes may thus act in interkingdom-signaling, a hypothesis that has recent support based on the effect of oxidized and epimerized bile acids on the function of regulatory T cells.^68, 69^

The concerted action of pairs of HSDHs result in bile acid products with reduced toxicity for microbes expressing the HSDH(s) or for an important inter-species partner, which was likely a factor in the evolution of these enzymes. Examples of strains of species capable of epimerizing bile acid hydroxyl groups are found in the literature, and the physicochemical properties and reduced toxicity of β-hydroxy bile acids are known, providing hypotheses for physiological function. *Clostridium limosum* (now *Hathewaya limosa*) expresses both bile acid-inducible NADP-dependent 7α- and 7β-HSDH capable of converting CDCA to UDCA.^70^ CDCA is more hydrophilic and more toxic to bacteria than UDCA.^6, 71^ Indeed, treatment with UDCA increases the hydrophilicity of the biliary pool, reducing cellular toxicity and improving biliary disorders.^72^ Similarly, strains of *Eggerthella lenta*^15, 41^ and *Ruminococcus gnavus*^41^ express both NADPH-dependent 3α- and 3β-HSDHs capable of forming 3β-bile acids (iso-bile acids). Iso-bile acids are also more hydrophilic and less toxic to bacteria than the α-hydroxy isomers.^41^ At least some strains of *R. gnavus* also express NADPH-dependent 7β-HSDH, contributing to the epimerization of CDCA to UDCA.^39^ It may be speculated that *R. gnavus* HSDHs function in detoxification of hydrophobic bile acids such as CDCA and DCA; however, further work is needed. Analogous to *E. lenta* and *R. gnavus, C. paraputrificum* is another example of a strain encoding multiple HSDHs that favor formation of β-hydroxy bile acids.^19^ *C. paraputrificum* strains encode the iso-bile acid pathway as well as NADPH-dependent 12β-HSDH.^19, 20^ While little is known about the biological effects of 12β-bile acids (epi-bile acids), the physicochemical properties relative to 12α-hydroxy bile acids should approximate that of iso- and urso-derivatives.^6, 41, 71^ An important question emerging from these observations is whether one particular epimeric product rather than another has important consequences on the fitness of the bacterium generating them, or if the increased hydrophilicity and reduced toxicity are the key factors.

Since the initial detection of epi-bile acids by Eneroth et. al. and Ali et. al.^16–18^, the measurement of bile acid metabolomes in clinical samples has become commonplace^73^, yet few studies measure or report epi-bile acids. Recently, 12β-hydroxy and 12-oxo-bile acids have been quantified in human feces by Franco et. al. (2019). 12-oxoLCA was the most abundant oxo-bile acid in feces at concentrations of about one half that of DCA in stool. While epiDCA itself was not measured, 3-oxo-12β-hydroxy-CDCA was shown at 12 ± 4 µg/g wet feces.^74^ Additionally, epiDCA has been reported in biliary bile of angelfish, likely produced from bacterial origin, so the 12β-HSDH gene may be widespread among resident microbiota of diverse vertebrate taxa.^75^ A critical limitation to the study of epi-bile acids is the absence of commercially available standards, although there are methods available for chemical synthesis.^76, 77^ The newly identified bile acid 12β-HSDHs could be employed for the enzymatic production of epi-bile acid standards from oxo-intermediates.

The physiological effects of epi-bile acids are poorly characterized, particularly in the GI tract. Borgstrӧm and colleagues compared infusion of CA, ursoCA, and epiCA on bile flow, lipid secretion, bile acid synthesis, and bile micellar formation. In contrast to ursoCA and CA, epiCA was secreted into bile in an unconjugated form. The 12β-hydroxyl group may hinder the enzyme responsible for conjugation. Additionally, epiCA infusion increased the rate of secretion of newly synthesized bile salts.^78^ Another study reported increased 12-oxoLCA levels in rats with high tumor incidence when they were fed a high safflower oil or corn oil diet.(Reddy 1984) While the toxicity of epi-bile acids has not yet been tested relative to the secondary bile acids DCA or LCA, both 12-oxoLCA and epiDCA are less hydrophobic than DCA by LC-MS (**Figures 2 & 3**). Due to the involvement of DCA in cancers of the liver and colon^7, 8^, bile acid 12β-HSDH may be of therapeutic importance in modulating the bile acid pool in favor of epiDCA over toxic DCA. Future studies with animal models will be imperative to determine the effects of epi-bile acids on host physiology.

## Materials and methods

### Bacterial strains and chemicals

*Clostridium paraputrificum* ATCC 25780 and *Clostridium scindens* ATCC 35704 were obtained from 80°C glycerol stocks from culture collections at the University of Illinois Urbana-Champaign (UIUC). *E. coli* DH5α (Turbo) competent cells from New England Biolabs (Ipswich, MA) and NovaBlue GigaSingles™ Competent cells from Novagen (San Diego, CA, USA) were used for cloning, and *E. coli* BL21-Codon-Plus (DE3) RIPL was purchased from Stratagene (La Jolla, CA, USA) and used for protein overexpression. 5β-Cholanic acid-3α, 7α, 12α-triol (CA), 5β-cholanic acid-3α,12α-diol (DCA), and 5β-cholanic acid-3α,7α-diol (CDCA) were purchased from Sigma-Aldrich (St. Louis, MO, USA). Authentic 5β-cholanic acid-3α,12β-diol (epiDCA) and 5β-cholanic acid-3α,7α,12β-diol (epiCA) were generously obtained from Lee R. Hagey (University of California, San Diego). Other bile acids were purchased from Steraloids (Newport, RI, USA). All other reagents were of the highest possible purity and purchased from Fisher Scientific (Pittsburgh, PA, USA).

### Whole cell bile acid conversion assay

*C. paraputrificum* ATCC 25780 and *C. scindens* ATCC 35704 were cultivated in anaerobic brain heart infusion (BHI) broth for 24 hrs. Two mL anaerobic BHI was inoculated with 1:10 dilution of either organism along with 50 μM bile acid substrate and incubated at 37°C for 24 hours. The bacterial cultures were centrifuged at 10,000 ⨉ g for 5 min to remove bacterial cells and the conditioned medium was adjusted to pH 3.0. Solid phase extraction was used to extract bile acid products from bacterial culture. Waters tC18 vacuum cartridges (3 cc) (Milford, MA, USA) were preconditioned with 6 mL 100% hexanes, 3 mL 100% acetone, 6 mL 100% methanol, and 6 mL water (pH 3.0). The conditioned medium was added to the cartridges and vacuum was applied to pull media through dropwise. Cartridges were washed with 6 mL water (pH 3.0) and 40% methanol. Bile acid products were eluted with 3 mL 100% methanol. Eluates were then evaporated under nitrogen gas and the residues dissolved in 200 μL 100% methanol for LC-MS analysis.

### Liquid chromatography-mass spectrometry

LC-MS analysis for all samples was performed using a Waters Acquity UPLC system coupled to a Waters SYNAPT G2-Si ESI mass spectrometer (Milford, MA, USA). LC was performed with a Waters Acquity UPLC HSS T3 C18 column (1.8 μm particle size, 2.1 mm x 100 mm) at a column temperature of 40°C. Samples were injected at 1 μL. Mobile phase A was water and B was acetonitrile. The mobile phase gradient was as follows: 0 min 100% mobile phase A, 0.5 min 100% A, 25 min 0% A, 25.1 min 100% A, 28 min 100% A. The flow rate was 0.5 mL/min.

MS was carried out in negative ion mode with a desolvation temperature of 300°C and desolvation gas flow of 700 L/hr. The capillary voltage was 3,000 V. Source temperature was 100°C and cone voltage was 30 V. Chromatographs and mass spectrometry data were analyzed using Waters MassLynx software (Milford, MA, USA).

### Isolation of genomic DNA

Genomic DNA was extracted from *C. paraputrificum* ATCC 25780 using the Fast DNA isolation kit from Mo-Bio (Carlsbad, CA, USA) according to the manufacturer’s protocol for polymerase chain reaction and molecular cloning applications.

### Heterologous expression of potential 12β-HSDH proteins

The pET-28a(+) and pET-46 Ek/LIC vectors were obtained from Novagen (San Diego, CA, USA). Restriction enzymes were purchased from NEB (Ipswich, MA, USA). Inserts were generated by PCR amplification with cloning primers from Integrative DNA Technologies (Coralville, IA, USA) of *C. paraputrificum* ATCC 25780 genomic DNA or genes synthesized in *E. coli* K12 codon usage (IDT, Coralville, IA, USA). Cloning primers and genes created by gene synthesis are listed in **Table S1**. Inserts were amplified using the Phusion High Fidelity Polymerase (Stratagene, La Jolla, CA, USA) and cloned into pET-28a(+) after insert and vector were double digested with the appropriate restriction endonuclease and treated with DNA Ligase, or annealed into pET-46 Ek/LIC after treatment with T4 DNA Polymerase. Recombinant plasmids were transformed via heat shock method, plated, and grown overnight at 37°C on lysogeny broth (LB) agar plates supplemented with antibiotic (50 µg/mL kanamycin or 100 µg/mL ampicillin). Vectors were either transformed into chemically competent *E. coli* DH5α cells and grown with kanamycin (pET-28a(+)) or transformed into NovaBlue GigaSingles™ Competent cells and grown with ampicillin (pET-46 Ek/LIC). A single colony from each transformation was inoculated into LB medium (5 mL) containing the corresponding antibiotic and grown to saturation. Recombinant plasmids were extracted from cell pellets using the QIAprep Spin Miniprep kit (Qiagen, Valencia, CA, USA). The sequence of the inserts was confirmed by Sanger sequencing (W. M. Keck Center for Comparative and Functional Genomics at the University of Illinois at Urbana-Champaign).

For protein expression, the extracted recombinant plasmids were transformed into *E. coli* BL-21 CodonPlus (DE3) RIPL chemically competent cells by heat shock method and cultured overnight at 37°C on LB agar plates supplemented with ampicillin or kanamycin (100 µg/ml; 50 µg/mL) and chloramphenicol (50 µg/ml). Selected colonies were inoculated into 10 mL of LB medium supplemented with antibiotics and grown at 37°C for 6 hours with vigorous aeration. The pre-cultures were added to fresh LB medium (1 L), supplemented with antibiotics, and aerated at 37°C until reaching an OD600nm of 0.3. IPTG was added to each culture at a final concentration of 0.1 mM to induce and the temperature was decreased to 16°C for a 16-hour incubation. Cells were pelleted and resuspended in binding buffer (20 mM Tris-HCl, 300 mM NaCl, 10 mM 2-mercaptoethanol, pH 7.9). The cells were subjected to five passages through an EmulsiFlex C-3 cell homogenizer (Avestin, Ottawa, Canada), and the cell debris was separated by centrifugation. The recombinant protein in the soluble fraction was then purified using TALON® Metal Affinity Resin (Clontech Laboratories, Mountain View, CA, USA) per the manufacturer’s protocol. The recombinant protein was eluted using an elution buffer composed of 20 mM Tris-HCl, 300 mM NaCl, 10 mM 2-mercaptoethanol, and 250 mM imidazole at pH 7.9. The resulting purified protein was analyzed using sodium dodecyl sulfate-polyacrylamide gel electrophoresis (SDS-PAGE). The observed subunit mass for each was calculated by migration distance of purified protein to standard proteins in ImageJ (https://imagej.nih.gov/ij/docs/faqs.html). TMHMM v. 2.0 was used to predict transmembrane helices.^22^

### Enzyme Assays

Pure recombinant 12β-HSDH reaction mixtures were made using 50 μM substrate, 150 μM cofactor and 10 nM enzyme in 150 mM NaCl, 50 mM sodium phosphate buffer at the pH optima 7.0 or 7.5. Reactions were monitored by spectrophotometric assay measuring the oxidation or reduction of NADP(H) aerobically at 340 nm (6,220 M^-1^.cm^-1^) continuously for 1.5 min on a NanoDrop 2000c UV-Vis spectrophotometer (Fisher Scientific, Pittsburgh, PA, USA) using a 10 mm quartz cuvette (Starna Cells, Atascadera, CA, USA). Additional reactions were incubated overnight at room temperature and extracted by vortexing with two volumes ethyl acetate twice. The organic layer was recovered and evaporated under nitrogen gas. The products were dissolved in 50 μL methanol and LC-MS was performed as described above or used for thin layer chromatography.

The buffers for investigation of the optimal pH of recombinant 12β-HSDH contained 150 mM NaCl and one of the following buffering agents: 50 mM sodium acetate (pH 6.0), 50 mM sodium phosphate (pH 6.5 to 7.5), and 50 mM Tris-Cl (pH 8.0). Substrate specificity was performed according to the above reaction conditions at the optimal pH.

The reaction mixtures for kinetic analysis were 10 nM enzyme, sodium phosphate buffer (pH 7.0), and 150 µM NADPH for varying concentrations of 12-oxoLCA or 80 µM 12-oxoLCA for varying NADPH concentrations in the reductive direction. The oxidative reaction mixture contained 10 nM enzyme, sodium phosphate buffer (pH 7.5), and 300 µM NADP^+^ when epiDCA concentrations were changed or 100 µM epiDCA when NADP^+^ was varied. Kinetic parameters were estimated with GraphPad Prism (GraphPad Prism, La Jolla, CA, USA) to fit the data using nonlinear regression to the Michaelis-Menten equation.

### Thin layer chromatography

Reaction mixtures were made using 50 μM substrate, 150 μM cofactor and 10 nM enzyme in 150 mM NaCl, 50 mM sodium phosphate buffer at pH 7.0. Reactions were incubated overnight at room temperature and extracted by vortexing with two volumes ethyl acetate twice. The organic layer was recovered and evaporated under nitrogen gas. The products were dissolved in 50 μL methanol and spotted on a TLC plate (silica gel IB2-F flexible TLC sheet, 20 x 20 cm, 250 μm analytical layer; J. T. Baker, Avantor Performance Materials, LLC, PA, USA). The steroids were separated with a 70:20:2 toluene–1,4-dioxane–acetic acid mobile phase and visualized using a 10% phosphomolybdic acid in ethanol spray and heating for 15 min at 100°C.^79^

### Native molecular weight determination

Size-exclusion chromatography was performed using a Superose 6 10/300 GL analytical column (GE Healthcare, Piscataway, NJ, USA) connected to an ÄKTAxpress chromatography system (GE Healthcare, Piscataway, NJ, USA) at 4°C. The column was equilibrated with 50 mM Tris-Cl and 150 mM NaCl at a pH of 7.5. The purified protein was loaded onto the analytical column at a concentration of 10 mg/mL and eluted at a flow rate of 0.3 ml/min. The native molecular mass of 12β-HSDH was determined by comparing its elution volume to that of Gel Filtration Standard proteins (Bio-Rad, Hercules, CA, USA): thyroglobulin, γ-globulin, ovalbumin, myoglobin, vitamin B12.

### Phylogenetic Analysis

The sequence of the *C. paraputrificum* 12β-HSDH protein (accession number WP_027099077.1) was used as query for a similarity search against the NCBI non-redundant protein database by BLASTP^80^, with a maximum E-value threshold of 1e-10 and a limit of 5,000 results. Retrieved sequences were aligned with Muscle v. 3.8.1551^81^ and analyzed by maximum likelihood with RAxML v. 8.2.11.^82^ Selection of the best-fitting amino acid substitution model and number of bootstrap pseudoreplicates were performed automatically, and substitution rate heterogeneity was modeled with gamma distributed rate categories. The resulting phylogenetic tree was formatted by Dendroscope v. 3.5.10^83^ and further cosmetic modifications were performed with the vector editor Inkscape, v. 0.92.4 (https://inkscape.org).

For closer analysis of the phylogenetic affiliation of *C. paraputrificum* ATCC 25780 12β-HSDH, sequences from the well-supported subtree where this sequence is located in the 5,000-sequence tree, plus an outgroup, were reanalyzed for confirming the relative placement of all sequences nearest to Cp12β-HSDH. The methods used were the same as described above for the full tree.

A maximum-likelihood tree of representative HSDH sequences was inferred by selecting sequences from each HSDH subfamily, based on the tree from Mythen et al. (2018)^15^, with the addition of eukaryotic, archaeal, and other bacterial sequences deposited in the public databases. Phylogenetic inference methods were the same as described above.

### Hidden Markov Model Search

A Hidden Markov Model (HMM) search was performed using a custom HMM profile against a concatenated file of metagenome assembled genomes (MAGs) from four publicly available cohorts.^30–33^ MAGs were filtered for genome completeness, quality, and contamination as described.^84^ For generation of the custom 12β-HSDH profile, reference sequences from the 12β-HSDHs characterized in this paper were aligned with MAFFT, manually trimmed, and constructed using hmmscan.^85^ The MAG database was searched using HMMSearch version 3.3.0^85^, using an individually identified cut-off of 350.00. Resulting hits were then filtered to remove results less than 70% completeness and closest matched species were recorded. The HMM search file is publicly available at: https://github.com/AnantharamanLab/doden_et_al_2021.

## Acknowledgments

We gratefully acknowledge support to J.M.R. for this work by the National Cancer Institute grant 1RO1 CA204808-01, as well as USDA Hatch ILLU-538-916 and Illinois Campus Research Board RB18068. H.L.D. is supported by the David H. and Norraine A. Baker Graduate Fellowship in Animal Sciences. We would like to express our very great appreciation to Dr. Lee R. Hagey for providing the critical substrates epiDCA and epiCA.

## References

1. Ridlon JM, Kang D-J, Hylemon PB. Bile salt biotransformations by human intestinal bacteria. J Lipid Res 2006; 47:241–59.

2. Vlahcevic ZR, Heuman DM, Hylemon PB. Physiology and pathophysiology of enterohepatic circulation of bile acids. In: Zakim D, Boyer T, editors. Hepatology: A Textbook of Liver Disease. Philadelphia: Saunders; 1996. page 376–417.

3. Dawson PA, Karpen SJ. Intestinal transport and metabolism of bile acids. J Lipid Res 2015; 56:1085–99.

4. Jones B V., Begley M, Hill C, Gahan CGM, Marchesi JR. Functional and comparative metagenomic analysis of bile salt hydrolase activity in the human gut microbiome. Proc Natl Acad Sci U S A 2008; 105:13580–5.

5. Ridlon JM, Harris SC, Bhowmik S, Kang DJ, Hylemon PB. Consequences of bile salt biotransformations by intestinal bacteria. Gut Microbes 2016; 7:22–39.

6. Watanabe M, Fukiya S, Yokota A. Comprehensive evaluation of the bactericidal activities of free bile acids in the large intestine of humans and rodents. J Lipid Res 2017; 58:1143– 52.

7. Bernstein C, Holubec H, Bhattacharyya AK, Nguyen H, Payne CM, Zaitlin B, Bernstein H. Carcinogenicity of deoxycholate, a secondary bile acid. Arch Toxicol 2011; 85:863– 71.

8. Yoshimoto S, Loo TM, Atarashi K, Kanda H, Sato S, Oyadomari S, Iwakura Y, Oshima K, Morita H, Hattori M, et al. Obesity-induced gut microbial metabolite promotes liver cancer through senescence secretome. Nature 2013; 499:97–101.

9. Wu JT, Gong J, Geng J, Song YX. Deoxycholic acid induces the overexpression of intestinal mucin, MUC2, via NF-kB signaling pathway in human esophageal adenocarcinoma cells. BMC Cancer 2008; 8:1–10.

10. Doden H, Sallam LA, Devendran S, Ly L, Doden G, Daniel SL, Ridlon JM. Metabolism of oxo-bile acids and characterization of recombinant 12α-hydroxysteroid dehydrogenases from bile acid 7α-dehydroxylating human gut bacteria. Appl Environ Microbiol 2018; 84:e00235–18.

11. Macdonald IA, Jellett JF, Mahony DE. 12alpha-Hydroxysteroid dehydrogenase from *Clostridium* group P strain C48-50 ATCC #29733: partial purification and characterization. J Lipid Res 1979; 20:234–9.

12. Harris JN, Hylemon PB. Partial purification and characterization of NADP-dependent 12α-hydroxysteroid dehydrogenase from *Clostridium leptum*. Biochim Biophys Acta 1978; 528:148–57.

13. Macdonald IA, Jellett JF, Mahony DE, Holdeman L V. Bile salt 3α- and 12α-hydroxysteroid dehydrogenases from *Eubacterium lentum* and related organisms. Appl Environ Microbiol 1979; 37:992–1000.

14. Macdonald IA, Meier EC, Mahony DE, Costain GA. 3α-, 7α- And 12α-hydroxysteroid dehydrogenase activities from Clostridium perfringens. Biochim Biophys Acta 1976; 450:142–53.

15. Mythen SM, Devendran S, Méndez-García C, Cann I, Ridlon JM. Targeted synthesis and characterization of a gene cluster encoding NAD(P)H-dependent 3α-, 3β-, and 12α-hydroxysteroid dehydrogenases from *Eggerthella* CAG:298, a gut metagenomic sequence. Appl Environ Microbiol 2018; 84:e02475–17.

16. Ali SS, Kuksis A, Beveridge JM. Excretion of bile acids by three men on corn oil and butterfat diets. Can J Biochem 1966; 44:1377–88.

17. Ali SS, Kuksis A, Beveridge JM. Excretion of bile acids by three men on a fat-free diet. Can J Biochem 1966; 44:957–69.

18. Eneroth P, Gordon B, Ryhage R, Sjövall J. Identification of mono- and dihydroxy bile acids in human feces by gas-liquid chromatography and mass spectrometry. J Lipid Res 1966; 7:511–23.

19. Edenharder R, Schneider J. 12β-Dehydrogenation of bile acids by *Clostridium paraputrificum*, *C. tertium*, and *C. difficile* and epimerization at carbon-12 of deoxycholic acid by cocultivation with 12α-dehydrogenating *Eubacterium lentum*. Appl Environ Microbiol 1985; 49:964–8.

20. Edenharder R, Pfützner A. Characterization of NADP-dependent 12β-hydroxysteroid dehydrogenase from *Clostridium paraputrificum*. Biochim Biophys Acta 1988; 962:362– 70.

21. Penning TM. Human hydroxysteroid dehydrogenases and pre-receptor regulation: Insights into inhibitor design and evaluation. J Steroid Biochem Mol Biol 2011; 125:46–56.

22. Krogh A, Larsson B, Von Heijne G, Sonnhammer ELL. Predicting transmembrane protein topology with a hidden Markov model: Application to complete genomes. J Mol Biol 2001; 305:567–80.

23. Kiu R, Caim S, Alcon-Giner C, Belteki G, Clarke P, Pickard D, Dougan G, Hall LJ. Preterm infant-associated *Clostridium tertium*, *Clostridium cadaveris*, and *Clostridium paraputrificum* strains: Genomic and evolutionary insights. Genome Biol Evol 2017; 9:2707–14.

24. Muñoz M, Restrepo-Montoya D, Kumar N, Iraola G, Herrera G, Ríos-Chaparro DI, Díaz-Arévalo D, Patarroyo MA, Lawley TD, Ramírez JD. Comparative genomics identifies potential virulence factors in *Clostridium tertium* and *C. paraputrificum*. Virulence 2019; 10:657–76.

25. Liu L, Aigner A, Schmid RD. Identification, cloning, heterologous expression, and characterization of a NADPH-dependent 7β-hydroxysteroid dehydrogenase from *Collinsella aerofaciens*. Appl Microbiol Biotechnol 2011; 90:127–35.

26. Wegner K, Just S, Gau L, Mueller H, Gérard P, Lepage P, Clavel T, Rohn S. Rapid analysis of bile acids in different biological matrices using LC-ESI-MS/MS for the investigation of bile acid transformation by mammalian gut bacteria. Anal Bioanal Chem 2017; 409:1231–45.

27. Sohn JH, Kwon KK, Kang JH, Jung HB, Kim SJ. *Novosphingobium pentaromativorans* sp. nov., a high-molecular-mass polycyclic aromatic hydrocarbon-degrading bacterium isolated from estuarine sediment. Int J Syst Evol Microbiol 2004; 54:1483–7.

28. Hashimoto T, Onda K, Morita T, Luxmy BS, Tada K, Miya A, Murakami T. Contribution of the estrogen-degrading bacterium *Novosphingobium* sp. strain JEM-1 to estrogen removal in wastewater treatment. J Environ Eng 2010; 136:890–6.

29. Gan HM, Hudson AO, Rahman AYA, Chan KG, Savka MA. Comparative genomic analysis of six bacteria belonging to the genus *Novosphingobium*: Insights into marine adaptation, cell-cell signaling and bioremediation. BMC Genomics 2013; 14:431.

30. Yu J, Feng Q, Wong SH, Zhang D, Yi Liang Q, Qin Y, Tang L, Zhao H, Stenvang J, Li Y, et al. Metagenomic analysis of faecal microbiome as a tool towards targeted non-invasive biomarkers for colorectal cancer. Gut 2017; 66:70–8.

31. Zeller G, Tap J, Voigt AY, Sunagawa S, Kultima JR, Costea PI, Amiot A, Böhm J, Brunetti F, Habermann N, et al. Potential of fecal microbiota for early-stage detection of colorectal cancer. Mol Syst Biol 2014; 10:1–18.

32. Vogtmann E, Hua X, Zeller G, Sunagawa S, Voigt AY, Hercog R, Goedert JJ, Shi J, Bork P, Sinha R. Colorectal cancer and the human gut microbiome: Reproducibility with whole-genome shotgun sequencing. PLoS One 2016; 11:e0155362.

33. Feng Q, Liang S, Jia H, Stadlmayr A, Tang L, Lan Z, Zhang D, Xia H, Xu X, Jie Z, et al. Gut microbiome development along the colorectal adenoma-carcinoma sequence. Nat Commun 2015; 6:1–13.

34. Chung WSF, Meijerink M, Zeuner B, Holck J, Louis P, Meyer AS, Wells JM, Flint HJ, Duncan SH. Prebiotic potential of pectin and pectic oligosaccharides to promote anti-inflammatory commensal bacteria in the human colon. FEMS Microbiol Ecol 2017; 93:1– 9.

35. Liu S, Zhao W, Liu X, Cheng L. Metagenomic analysis of the gut microbiome in atherosclerosis patients identify cross-cohort microbial signatures and potential therapeutic target. FASEB J 2020; 34:14166–81.

36. Aigner A, Gross R, Schmid R, Braun M, Mauer S. Novel 12α-hydroxysteroid dehydrogenases, production and use therof. 2011; :US patent 20110091921A1.

37. Baron SF, Franklund C V., Hylemon PB. Cloning, sequencing, and expression of the gene coding for bile acid 7α-hydroxysteroid dehydrogenase from *Eubacterium* sp. strain VPI 12708. J Bacteriol 1991; 173:4558–69.

38. Yoshimoto T, Higashi H, Kanatani A, Lin XS, Nagai H, Oyama H, Kurazono K, Tsuru D. Cloning and sequencing of the 7α-hydroxysteroid dehydrogenase gene from *Escherichia coli* HB101 and characterization of the expressed enzyme. J Bacteriol 1991; 173:2173–9.

39. Lee JY, Arai H, Nakamura Y, Fukiya S, Wada M, Yokota A. Contribution of the 7β-hydroxysteroid dehydrogenase from *Ruminococcus gnavus* N53 to ursodeoxycholic acid formation in the human colon. J Lipid Res 2013; 54:3062–9.

40. Ferrandi EE, Bertolesi GM, Polentini F, Negri A, Riva S, Monti D. In search of sustainable chemical processes: Cloning, recombinant expression, and functional characterization of the 7α- and 7β-hydroxysteroid dehydrogenases from *Clostridium absonum*. Appl Microbiol Biotechnol 2012; 95:1221–33.

41. Devlin AS, Fischbach MA. A biosynthetic pathway for a prominent class of microbiota-derived bile acids. Nat Chem Biol 2015; 11:685–90.

42. Doden HL, Pollet RM, Mythen SM, Wawrzak Z, Devendran S, Cann I, Koropatkin NM, Ridlon JM. Structural and biochemical characterization of 20β-hydroxysteroid dehydrogenase from *Bifidobacterium adolescentis* strain L2-32. J Biol Chem 2019; 294:12040–53.

43. Devendran S, Méndez-García C, Ridlon JM. Identification and characterization of a 20β-HSDH from the anaerobic gut bacterium *Butyricicoccus desmolans* ATCC 43058. J Lipid Res 2017; 58:916–25.

44. Ridlon JM, Ikegawa S, Alves JMP, Zhou B, Kobayashi A, Iida T, Mitamura K, Tanabe G, Serrano M, De Guzman A, et al. *Clostridium scindens*: a human gut microbe with a high potential to convert glucocorticoids into androgens. J Lipid Res 2013; 54:2437–49.

45. Bernardi R, Doden H, Melo M, Devendran S, Pollet R, Mythen S, Bhowmik S, Lesley S, Cann I, Luthey-Schulten Z, et al. Bacteria on steroids: the enzymatic mechanism of an NADH-dependent dehydrogenase that regulates the conversion of cortisol to androgen in the gut microbiome. 2020; :bioRxiv 2020.06.12.149468. Available from: https://doi.org/10.1101/2020.06.12.149468

46. Fahrbach M, Kuever J, Meinke R, Kämpfer P, Hollender J. *Denitratisoma oestradiolicum* gen. nov., sp. nov., a 17β-oestradiol-degrading, denitrifying betaproteobacterium. Int J Syst Evol Microbiol 2006; 56:1547–52.

47. Ricaboni D, Mailhe M, Vitton V, Cadoret F, Fournier PE, Raoult D. *‘Intestinibacillus massiliensis’* gen. nov., sp. nov., isolated from human left colon. New Microbes New Infect 2017; 17:18–20.

48. Morris DJ, Ridlon JM. Glucocorticoids and gut bacteria: “The GALF Hypothesis” in the metagenomic era. Steroids 2017; 125:1–13.

49. Sherrod JA, Hylemon PB. Partial purification and characterization of NAD-dependent 7α-hydroxysteroid dehydrogenase from Bacteroides thetaiotaomicron. 1977; 486:351–8.

50. Edenharder R, Mielek K. Epimerization, oxidation and reduction of bile acids by *Eubacterium lentum*. Syst Appl Microbiol 1984; 5:287–98.

51. Snyder ML. The serologic agglutinatin of the obligate anaerobes *Clostridium paraputrificum* (Beinstock) and *Clostridium capitovalis* (Snyder and Hall). J Bacteriol 1936; 32:401–10.

52. Harris SC, Devendran S, Méndez-García C, Mythen SM, Wright CL, Fields CJ, Hernandez AG, Cann I, Hylemon PB, Ridlon JM. Bile acid oxidation by *Eggerthella lenta* strains C592 and DSM 2243. Gut Microbes 2018; 9:523–39.

53. Filling C, Berndt KD, Benach J, Knapp S, Prozorovski T, Nordling E, Ladenstein R, Jörnvall H, Oppermann U. Critical residues for structure and catalysis in short-chain dehydrogenases/reductases. J Biol Chem 2002; 277:25677–84.

54. Grimm C, Maseri E, Möbusi E, Klebe G, Reuter K, Ficner R. The crystal structure of 3α-hydroxysteroid dehydrogenase/carbonyl reductase from *Comamonas testosteroni* shows a novel oligomerization pattern within the short chain dehydrogenase/reductase family. J Biol Chem 2000; 275:41333–9.

55. Savino S, Ferrandi EE, Forneris F, Rovida S, Riva S, Monti D, Mattevi A. Structural and biochemical insights into 7β-hydroxysteroid dehydrogenase stereoselectivity. Proteins 2016; 84:859–65.

56. Jang LG, Choi G, Kim SW, Kim BY, Lee S, Park H. The combination of sport and sport-specific diet is associated with characteristics of gut microbiota: An observational study. J Int Soc Sports Nutr 2019; 16:1–10.

57. Ma S, You Y, Huang L, Long S, Zhang J, Guo C, Zhang N, Wu X, Xiao Y, Tan H. Alterations in gut microbiota of gestational diabetes patients during the first trimester of pregnancy. Front Cell Infect Microbiol 2020; 10:1–14.

58. Amaruddin AI, Hamid F, Koopman JPR, Muhammad M, Brienen EAT, van Lieshout L, Geelen AR, Wahyuni S, Kuijper EJ, Sartono E, et al. The bacterial gut microbiota of schoolchildren from high and low socioeconomic status: A study in an urban area of Makassar, Indonesia. Microorganisms 2020; 8:1–12.

59. Doumatey AP, Adeyemo A, Zhou J, Lei L, Adebamowo SN, Adebamowo C, Rotimi CN. Gut microbiome profiles are associated with type 2 diabetes in urban africans. Front Cell Infect Microbiol 2020; 10:1–13.

60. Turpin W, Espin-Garcia O, Xu W, Silverberg MS, Kevans D, Smith MI, Guttman DS, Griffiths A, Panaccione R, Otley A, et al. Association of host genome with intestinal microbial composition in a large healthy cohort. Nat Genet 2016; 48:1413–7.

61. Labbé A, Ganopolsky JG, Martoni CJ, Prakash S, Jones ML. Bacterial bile metabolising gene abundance in Crohn’s, ulcerative colitis and type 2 diabetes metagenomes. PLoS One 2014; 9:e115175.

62. Lepercq P, Gérard P, Béguet F, Raibaud P, Grill JP, Relano P, Cayuela C, Juste C. Epimerization of chenodeoxycholic acid to ursodeoxycholic acid by *Clostridium baratii* isolated from human feces. FEMS Microbiol Lett 2004; 235:65–72.

63. Horinouchi M, Hayashi T, Koshino H, Malon M, Yamamoto T, Kudo T. Identification of genes involved in inversion of stereochemistry of a C-12 hydroxyl group in the catabolism of cholic acid by *Comamonas testosteroni* TA441. J Bacteriol 2008; 190:5545–54.

64. Holert J, Kulić Ž, Yücel O, Suvekbala V, Suter MJF, Möller HM, Philipp B. Degradation of the acyl side chain of the steroid compound cholate in *Pseudomonas* sp. strain Chol1 proceeds via an aldehyde intermediate. J Bacteriol 2013; 195:585–95.

65. Edenharder R, Pfützner M, Hammann R. NADP-dependent 3β-, 7α- and 7β-hydroxysteroid dehydrogenase activities from a lecithinase-lipase-negative *Clostridium* species 25.11.c. Biochim Biophys Acta 1989; 1002:37–44.

66. Ridlon JM, Devendran S, Alves JM, Doden H, Wolf PG, Pereira G V., Ly L, Volland A, Takei H, Nittono H, et al. The ‘in vivo lifestyle’ of bile acid 7α-dehydroxylating bacteria: comparative genomics, metatranscriptomic, and bile acid metabolomics analysis of a defined microbial community in gnotobiotic mice. Gut Microbes 2020; 11:381–404.

67. Wahlström A, Kovatcheva-Datchary P, Ståhlman M, Bäckhed F, Marschall HU. Crosstalk between bile acids and gut microbiota and its impact on farnesoid X receptor signalling. Dig Dis 2017; 35:246–50.

68. Song X, Sun X, Oh SF, Wu M, Zhang Y, Zheng W, Geva-Zatorsky N, Jupp R, Mathis D, Benoist C, et al. Microbial bile acid metabolites modulate gut RORγ+ regulatory T cell homeostasis. Nature 2020; 577:410–5.

69. Campbell C, McKenney PT, Konstantinovsky D, Isaeva OI, Schizas M, Verter J, Mai C, Jin WB, Guo CJ, Violante S, et al. Bacterial metabolism of bile acids promotes generation of peripheral regulatory T cells. Nature 2020; 581:475–9.

70. Sutherland JD, Williams CN. Bile acid induction of 7α- and 7β-hydroxysteroid dehydrogenases in *Clostridium limosum*. J Lipid Res 1985; 26:344–50.

71. Hofmann AF, Roda A. Physicochemical properties of bile acids and their relationship to biological properties: An overview of the problem. J Lipid Res 1984; 25:1477–89.

72. Goossens J, Bailly C. Ursodeoxycholic acid and cancer: From chemoprevention to chemotherapy. Pharmacol Ther 2019; 203:107396.

73. Liu Y, Rong Z, Xiang D, Zhang C, Liu D. Detection technologies and metabolic profiling of bile acids: A comprehensive review. Lipids Health Dis 2018; 17:121.

74. Franco P, Porru E, Fiori J, Gioiello A, Cerra B, Roda G, Caliceti C, Simoni P, Roda A. Identification and quantification of oxo-bile acids in human faeces with liquid chromatography–mass spectrometry: A potent tool for human gut acidic sterolbiome studies. J Chromatogr A 2019; 1585:70–81.

75. Hofmann AF, Hagey LR, Krasowski MD. Bile salts of vertebrates: Structural variation and possible evolutionary significance. J Lipid Res 2010; 51:226–46.

76. Chang FC. Potential bile acid metabolites. 5.1 12β-hydroxy acids by stereoselective reduction. Synth Commun 1981; 11:875–9.

77. Iida T, Momose T, Chang FC, Nambara T. Potential bile acid metabolites. XI. Syntheses of stereoisomeric 7,12-dihydroxy-5α-cholanic acids. Chem Pharm Bull 1986; 34:1934–8.

78. Borgström B, Barrowman J, Krabisch L, Lindström M, Lillienau J. Effects of cholic acid, 7β-hydroxy- and 12β-hydroxy-isocholic acid on bile flow, lipid secretion and bile acid synthesis in the rat. Scand J Clin Lab Invest 1986; 46:167–75.

79. Eneroth P. Thin-layer chromatography of bile acids. J Lipid Res 1963; 4:11–6.

80. Camacho C, Coulouris G, Avagyan V, Ma N, Papadopoulos J, Bealer K, Madden TL. BLAST+: Architecture and applications. BMC Bioinformatics 2009; 10:1–9.

81. Edgar RC. MUSCLE: Multiple sequence alignment with high accuracy and high throughput. Nucleic Acids Res 2004; 32:1792–7.

82. Stamatakis A. RAxML version 8: A tool for phylogenetic analysis and post-analysis of large phylogenies. Bioinformatics 2014; 30:1312–3.

83. Huson DH, Richter DC, Rausch C, Dezulian T, Franz M, Rupp R. Dendroscope: An interactive viewer for large phylogenetic trees. BMC Bioinformatics 2007; 8:460.

84. Pasolli E, Asnicar F, Manara S, Zolfo M, Karcher N, Armanini F, Beghini F, Manghi P, Tett A, Ghensi P, et al. Extensive unexplored human microbiome diversity revealed by over 150,000 genomes from metagenomes spanning age, geography, and lifestyle. Cell 2019; 176:649–62.

85. Eddy SR. Accelerated profile HMM searches. PLoS Comput Biol 2011; 7:e1002195.

