## Supplemental Material for "Completion of the gut microbial epi-bile acid pathway"

*[a]Carl R. Woese Institute for Genomic Biology, University of Illinois at Urbana Champaign, Urbana, IL, USA; [b]Department of Animal Sciences, University of Illinois at Urbana Champaign, Urbana, IL, USA; [c]Division of Nutritional Sciences, University of Illinois at Urbana Champaign, Urbana, IL, USA; [d]Institute for Health Research and Policy, University of Illinois, Chicago, IL, USA; [e]Cancer Education and Career Development Program, University of Illinois, Chicago, IL, USA; [f]Cancer Center at Illinois, Urbana, IL, USA; [g]Department of Bacteriology, University of Wisconsin–Madison, Madison, WI, USA; [h]Department of Parasitology, Institute of Biomedical Sciences, University of São Paulo, São Paulo, Brazil; [i]Department of Microbiology and Immunology, Virginia Commonwealth University, Richmond, VA, USA*

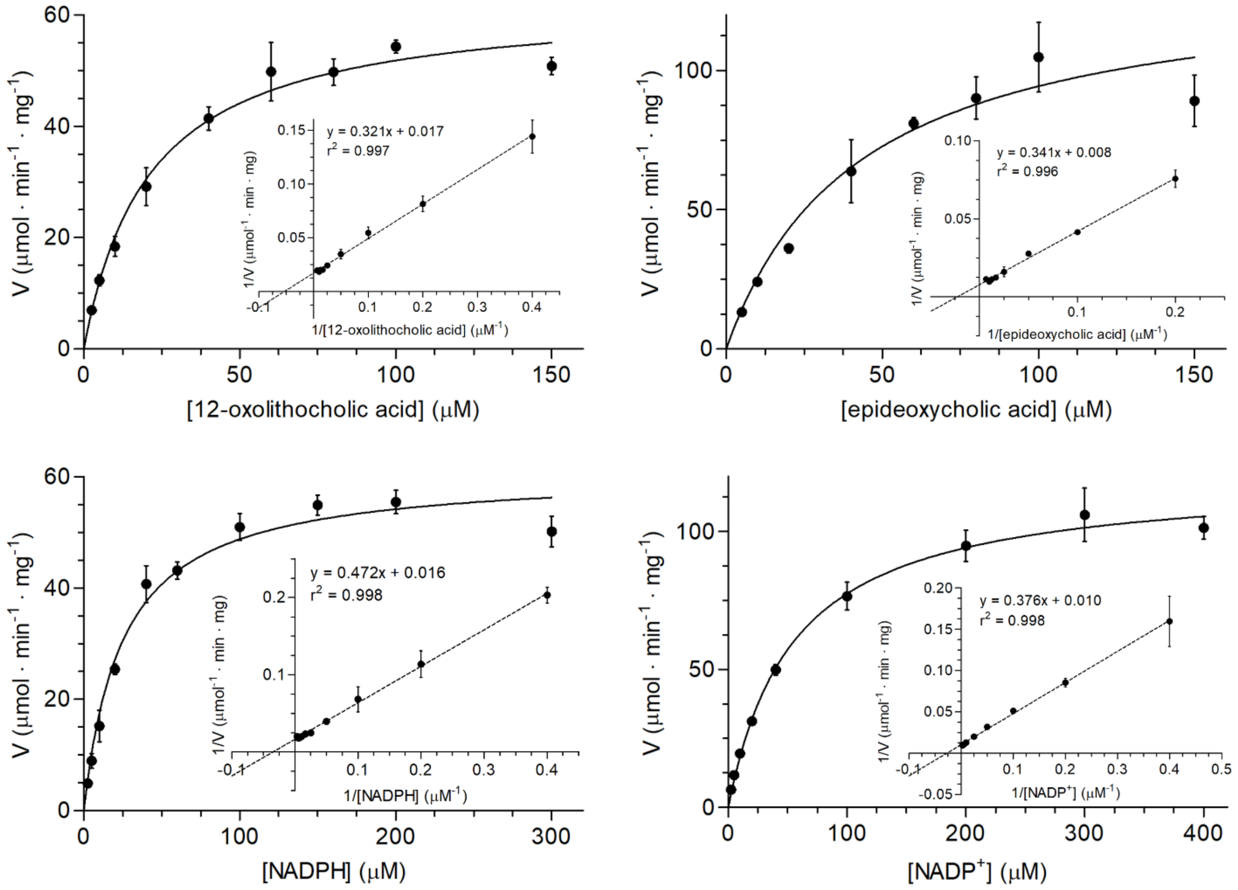

**Figure S1. Kinetic analysis of Cp12β-HSDH activity.** Michaelis-Menten and Lineweaver-Burk plots of initial enzyme velocity with varying concentration of substrate (top panels) and cofactor (bottom panels) in both the reductive (left panels) and oxidative (right panels) directions. See Materials and Methods for enzyme reactions. Oxidation/reduction of NAD(H) was measured by continuous spectrophotometry at 340 nm for 1.5 min and the initial velocities were used to calculate kinetic constants (Table 1). Each point represents the mean ± SD of three or more replicates.

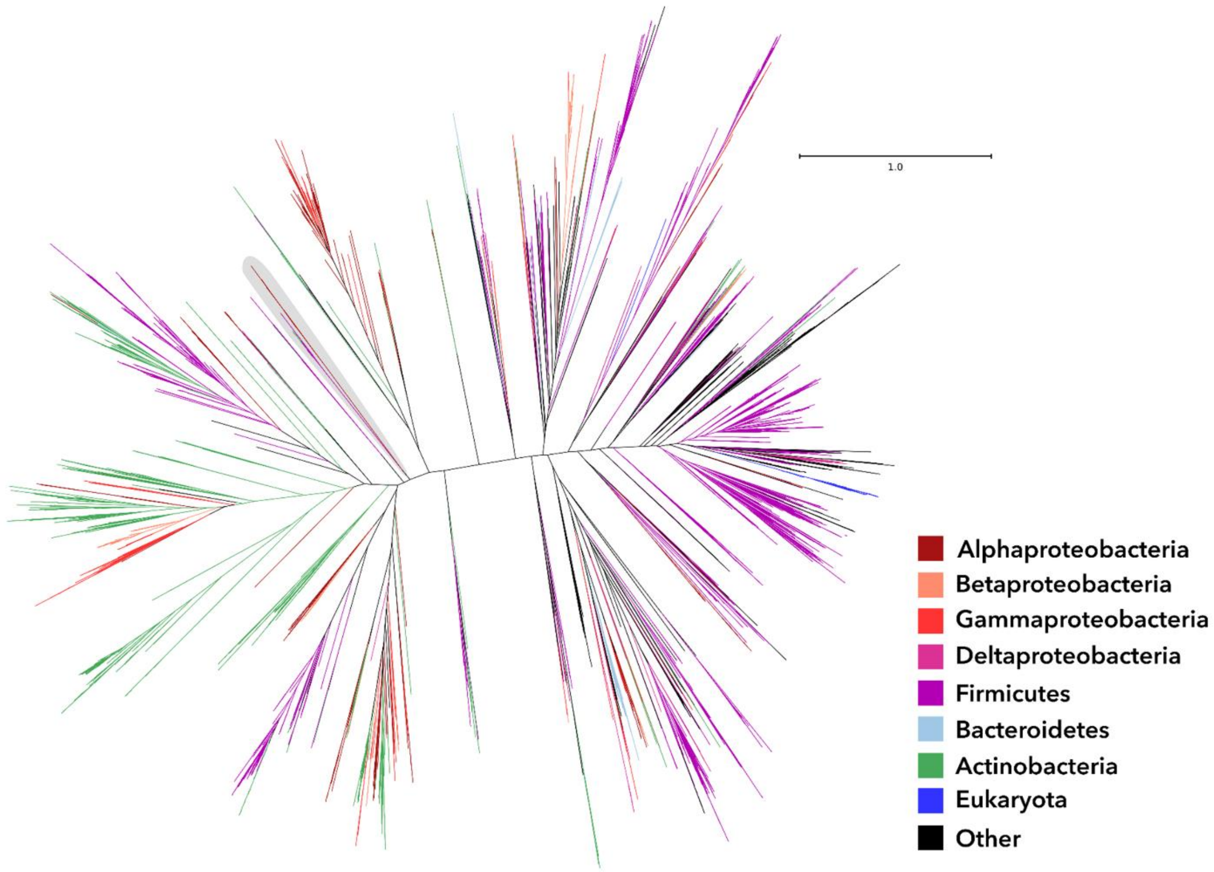

**Figure S2. 12β-HSDH 5,000-member tree.** Maximum likelihood phylogeny for the five thousand sequences from the NR database that are most similar to *C. paraputrificum* 12β-HSDH (WP_027099077.1). Taxonomic affiliations are indicated by branch colors as specified in the legend. The region highlighted in gray shows the sequences that were reanalyzed to generate the phylogeny shown in Figure 5.

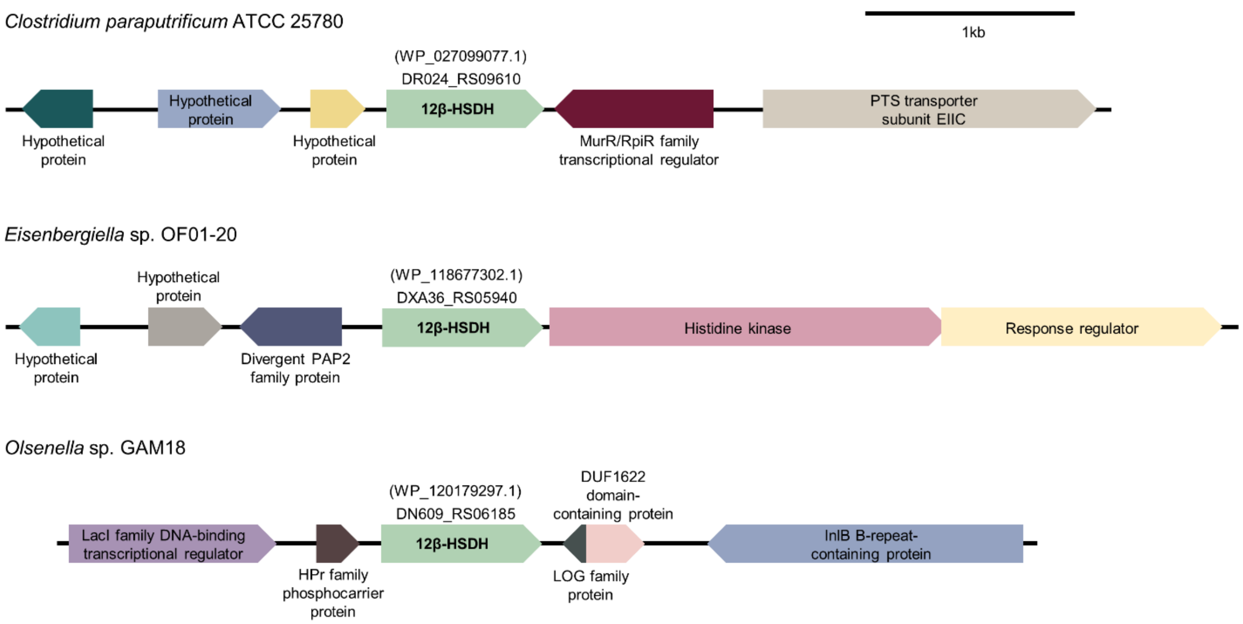

**Figure S3. Genomic context of 12β-HSDH genes from *Clostridium paraputrificum* ATCC 25780, *Eisenbergiella* sp. OF01-20, and *Olsenella* sp. GAM18.** Genomic context was identified using the NCBI nucleotide graphics function.

**Table S1.** Cloning primers for putative 12β-HSDHs and synthesized genes.

| Gene ID<br>Protein Accession | Primer pair | Vector | MW (kDa)[a] | Extinction coefficient ($M^{-1}$•$cm^{-1}$) |
|---|---|---|---|---|
| DR024_RS09610<br>WP_027099077.1 | ATATATCATATGATGAAAGAGTTAAATGAGAAAGTAGCTATTATTACAGG (5'-Forward Primer-3')<br>ATATATCTCGAGTTAAGGAGCTATTGAGTATCCACCATCC (5'-Reverse Primer-3') | pET-28a(+) | 27.35 | 31,775 |
| DR024_RS11955WP_027098355.1 | GACGACGACAAGATGATTGAAAAAACGTTTCCAAAAACTTTTCCTACT<br>GAGGAGAAGCCCGGTTTAACTATTAACTATTTCCCCTCCATTAACATGAAT | pET-46 Ek/LIC | 31.39 | 19,160 |
| DR024_RS09610<br>WP_027097937.1 | ATATATGGATCCATGTCAAAGTTATTAGGTAAGGTAGCCATTATTAC<br>ATATATCTCGAGTTAAATATACCCTCCATCCACTCTAAGTATCTG | pET-28a(+) | 26.26 | 14,565 |
| DR024_RS15375<br>WP_027098604.1 | GACGACGACAAGATGAAAAATAAATTCACACTAATAACTGGTGGAAGTGA<br>GAGGAGAAGCCCGGTCTAATACTTATTGCTTTTTAAGACCACTTTTCTACTTAATGCT | pET-46 Ek/LIC | 28.75 | 18,005 |
| DR024_RS02235<br>WP_027096909.1 | ATATATCATATGATGAAGCAAATAAAAATAGCTAATACAGATATGGAAGT<br>ATATATCTCGAGTTAAGGAAGTGTATTTCCTGCTGCCTTA | pET-28a(+) | 34.68 | 31,650 |
| DR024_RS08875<br>WP_027099631.1 | ATATATGCTAGCATGTCTGTATTGGAGAATAATTATAGGGATTTTAAATGT<br>ATATATCTCGAGTTAGCTATCTACCATAGTTCCACCATTAACA | pET-28a(+) | 32.86 | 34,060 |
| DXA36_RS05940<br>WP_118677302.1 | ATATATCATATGATGAAACAGTTGAATGAGAAAGTAGCCAT<br>ATATATCTCGAGTTATGGCATAATCGAATAGGCTCCATCAAC | pET-28a(+) | 27.34 | 33,390 |
| | ATGAAACAGTTGAATGAGAAAGTAGCCATCGTAACTGGTGCTGGTCAAGGCATTGGTCAAGGAATTGCATTA<br>TGTTTGGGGAAGCGCGGTGTTAAGGTAGTGTGCGTAGGTCGCCGTCCCGAACCTATTGAAGCTACTGCTAAA<br>GAAATTCGTGACTTAGGTGGTGAGTCTTTCGCTATGACCTGTGACACAGCGGATCGTGATCGTGTTAAAGAGG<br>TAGTCGCAAAAACCGTGGAAACGTACAAAACTGTAGATGTAATGATTAATAACGCGCAAAGTTTACCCGGAT<br>CCGCCCCTGTTGAGGAGGTCACTTATGAAATGATGTACACGGCATGGAGTACTGGAACACTTGGATCATTAA<br>ACTTTATGCAAGAATGTTTCCCTTACATGAAAGAGCAAGGGGAAGGGCGTGTAATCAACTTTGCGTCAGCCA<br>CTGGTATGTTCGGGTATGCTGGTAATCTTGCGTATGGTTGTAACAAAGAAGCCATTCGTGGCTTGACAAAGAT<br>TGCAGCGAAGGAGTGGGGCAAATACGGGATCTGTGTAAACTGTGTCCTGCCAGGCGCTGAGTCCCCAGCTGC<br>TAAGATCTGGGCTGAAAAATTTCCAGAAAAGTATGCGGAGATCTTGGAGCAACAACCTATGAAACGCTTAGG<br>AGACGCTGAAAAGGACATCGCGCCGGTGATTGCTTTCTTATCGGGTCCAGACAGCTGTTATTACAGTGGTCAA<br>TGCCTGTTGGTTGATGGAGCCTATTCGATTATGCCATAA[b] | | | |
| DN609_RS06185<br>WP_120179297.1 | ATATATCATATGATGAAACAACTTAATGAAAAAGTTGCAATTGTTACT<br>ATATATCTCGAGCTAGGGCATGATACTATTAGCTCCGTC | pET-28a(+) | 26.87 | 27,180 |
| | ATGAAACAACTTAATGAAAAAGTTGCAATTGTTACTGGTGCAGGGCAAGGGATCGGAAAAGGGATCGCCCTT<br>TGTTTAGCTAAACGTGGTGTAAAGGTCGTGTGCACGGGACGCCGTGAAGCACCAATCCAACAGACTGTAGCA<br>GAAATCGAAGAGTTAGGAGGTCAAGGACTTGCTATGACTTGTGACTCAGCAGACCGTGCGCGCGTCGAAGAG<br>GTGGTCAAGGCCGCGGTCGATACGTTTGGCTCCATCGATGTGATTGTAAATAATGGACAAGCCATTGTCCCCT<br>CTGCGCCTGTCGAGGACACTACATACGAGAACATGCTGGCGGCATGGCAAAGTGGCACTATCGGCTCGTTAA<br>ATTACATGCAGGCTGCATTTCCCCACATGAAGGAGCAGCACGAGGGGCGTATTATCAATTTCGCCTCCGCTAC<br>CGGAATGTTCGGCATTGCGGGACAACTGGCTTACGGCAGCAATAAAGAGGCACTTCGCGGTCTTACAAAGAT<br>CGCCGCTAAGGAATGGGGACAGTATGGAATTTGCGTTAACGTAGTATTGCCGGGGGCGGAATCCCCAGCCGC<br>CAAGGCGTGGGCCGAGAAATTTCCAGAAGAATACCAGAAACAGGTTATGTTAAATCCAATGCATCGTTTTGG<br>GGACCCAGAAGATGACATCGCGCCAGTTGTGGCATTCCTGGCGGGTCCTGATAGTTGCTATTACTCCGGCCAA<br>TCAGTCATCGTGGACGGAGCTAATAGTATCATGCCCTAG[b] | | | |
| PMI02_RS03035<br>WP_007678535.1 | ATATATGGATCCATGTCCAACGAGAGTCGTCGTTTAGA<br>ATATATCTCGAGTCAAAGCATTGTGGTGCCACCC | pET-28a(+) | 27.05 | 22,590 |
| | ATGTCCAACGAGAGTCGTCGTTTAGAAGGTAAAGTTGCAATTGTCACGGGGGCAGGTCAAGGGATCGGAGAG<br>GCTATCGCAGTTGGATACGCAGCTGCTGGAGCAAAAGTCTTGATCACGGGACGCACTCAGAGTAAGTTAGAT<br>GACGTTGTAAGTAAGATTGAAGCTGCCTCAGGGACAGCGATTGGGATGGAGGCATTGGCTGGTAACCGTGAA | | | |

CATGCTGCGCTTACAGTTGATCGCGCCATTTCAGAATGGGGGCGCGTCGACGTACTGGTTAATAATGCGCATA
CGTTTACGGATTTTTTGCCCATCGAGGACCCAAAGATGGAAGAGAATTGCAATACGGACATCCAATCAGCTTT
TTTTGGCAGTCTGCAGTTGATGCAATTGTGCCACCCCCATATGGCAGCCCAAGGTGGCGGCTCAATTATTAAC
ATGGGTACTGGTTTTTCTATTCGTTGCGAGCCGGGTTTTTTGGCCTACGCCGCTACTAAGGAAGCGATCCGCG
TATTGACCAAGACCGCAGCGAAGGAGTGGGGGAAACATAAGATCCGTGTGAATACGATCCTTCCTTCTGCTC
TGACGCCAAAAAGTATCTGGTATTTAGAGGACTCAGGAACATACGACCAGGAATTGGCTAAGGTGGCTATGG
GTCGTTTCGGCGAGCCGAAAGATATCGCGCCAACAGCGGTCTTTCTGGCCTCTGACGATTCAGATTTTGTGAC
GGGGCAGACTATCGGGGTGGAGGGTGGCACCACAATGCTTTGA[b]

[a]Deduced recombinant protein molecular weight.
[b]Gene synthesized by Integrated DNA Technologies with *E.coli* K12 codon optimization.

**Table S2:** Sequences used in hydroxysteroid dehydrogenase phylogenetic analysis.

| Accession No | Function | Organism | Reference |
|---|---|---|---|
| WP_027099077.1 | 12β-HSDH | *Clostridium paraputrificum* ATCC 25780 | Current study |
| WP_007678535.1 | putative 12β-HSDH | *Novoshingobium* sp. AP12 | Current study |
| WP_120179297.1 | 12β-HSDH | *Olsenella* sp. GAM18 | Current study |
| WP_009140706.1 | putative 12β-HSDH | *Collinsella tanakaei* | Current study |
| WP_117888595.1 | putative 12β-HSDH | *Ruminococcus* sp. AF14-10 | Current study |
| WP_005612898.1 | putative 12β-HSDH | *Ruminococcus lactaris* | Current study |
| WP_118677302.1 | 12β-HSDH | *Eisenbergiella* sp. OF01-20 | Current study |
| EDS06338.1 | 12α-HSDH | *Clostridium scindens* ATCC 35704 | Doden Appl. Environ. Microbiol. 2018 |
| EEG75500.1 | 12α-HSDH | *Clostridium hylemonae* DSM 15053 | Doden Appl. Environ. Microbiol. 2018 |
| EEA85268.1 | 12α-HSDH | *Peptacetobacter hiranonis/Clostridium hiranonis* DSM 13275 | Doden Appl. Environ. Microbiol. 2018 |
| CDD59475.1 | 12α-HSDH | *Eggerthella* sp. CAG:298 | Mythen Appl. Environ. Microbiol. 2018 |
| ERJ00208.1 | 12α-HSDH | *Clostridium* sp. strain ATCC 29733 | Macdonald J. Lipid Res. 1979; Aigner US patent 2011 |
| WP_096515955.1 | putative 12α-HSDH | *Clostridium perfringens* | Doden Appl. Environ. Microbiol. 2018 |
| WP_011406312.1 | putative 12α-HSDH | *Methanosphaera stadtmanae* DSM 3091 | Kisiela J. Steroid Biochem. Mol. Biol. 2012 |
| ABQ87936.1 | putative 12α-HSDH | *Methanobrevibacter smithii* ATCC 35061 | Kisiela J. Steroid Biochem. Mol. Biol. 2012 |
| EBA39192.1 | putative 12α-HSDH | *Collinsella aerofaciens* ATCC 25986 | Kisiela J. Steroid Biochem. Mol. Biol. 2012 |
| WP_006235414.1 | putative 12α-HSDH | *Collinsella aerofaciens* ATCC 25986 | Kisiela J. Steroid Biochem. Mol. Biol. 2012 |
| EEG90138.1 | putative 12α-HSDH | *Coprococcus comes* ATCC 27758 | Kisiela J. Ster. Biochem. Mol. Biol. 2012 |
| CDD59474.1 | 3α-HSDH | *Eggerthella* sp. CAG:298 | Mythen Appl. Environ. Microbiol. 2018 |
| ACV54671.1/ WP_009306474.1 | 3α-HSDH | *Eggerthella lenta* DSM 2243 | Devlin Nat. Chem. Biol. 2015 |
| EDN77529.1 | 3α-HSDH | *Ruminococcus gnavus* ATCC 29149 | Devlin Nat. Chem. Biol.2015 |
| P19337.1 | 3α-HSDH | *Clostridium scindens* ATCC 35704 | Mallonee Curr. Microbiol. 1995; Bhowmik Proteins 2014 |
| ACF20977.1 | 3α-HSDH | *Clostridium hylemonae* DSM 15053 | Ridlon Anaerobe 2010 |
| EEA86309.1 | 3α-HSDH | *Clostridium hiranonis* DSM 13275 | Wells Appl. Environ. Microbiol. 2000 |
| EEG48917.1 | putative 3α-HSDH | *Blautia hydrogenotrophica* DSM 10507 | Kisiela J. Steroid Biochem. Mol. Biol. 2012 |
| EDR48431.1 | putative 3α-HSDH | *Dorea formicigenerans* ATCC 27755 | Kisiela J. Steroid Biochem. Mol. Biol. 2012 |
| EDK24466.1 | putative 3α-HSDH | *Ruminococcus torques* ATCC 27756 | Kisiela J. Steroid Biochem. Mol. Biol. 2012 |
| EDM85978.1 | putative 3α-HSDH | *Ruminococcus obeum* ATCC 29174 | Kisiela J. Steroid Biochem. Mol. Biol. 2012 |
| EDT24590.1 | putative 3α-HSDH | *Clostridium perfringens* B str. ATCC 3626 | Macdonald Biochim. Biophys. Acta 1976; Hirano Appl. Environ. Microbiol. 1981 |
| CDD59473.1 | 3β-HSDH | *Eggerthella* sp. CAG:298 | Mythen Appl. Environ. Microbiol. 2018 |
| ACV55294.1 | 3β-HSDH | *Eggerthella lenta* DSM 2243 | Devlin Nat. Chem. Biol. 2015 |
| ACV54192.1 | 3β-HSDH | *Eggerthella lenta* DSM 2243 | Devlin Nat. Chem. Biol. 2015 |
| EDN78833.1 | 3β-HSDH | *Ruminococcus gnavus* ATCC 29149 | Devlin Nat. Chem. Biol. 2015 |
| BAA01384.1 | 7α-HSDH | *Escherichia coli* | Yoshimoto J. Bacteriol. 1991 |
| AAA53556.1 | 7α-HSDH | *Clostridium sordellii* ATCC 9714 | Coleman J. Bacteriol. 1994 |
| AAB61151.1 | 7α-HSDH | *Clostridium scindens* VPI 12708 | Baron J. Bacteriol. 1991 |
| WP_005792012.1 | 3-oxoacyl-[acyl-carrier-protein] reductase | *Bacteroides fragilis* ATCC 25285 | Doden Appl. Environ. Microbiol. 2018 |
| WP_006440226.1 | putative 7α-HSDH | *Clostridium hiranonis* DSM 13275/*Peptacetobacter hiranonis* | Kisiela J. Steroid Biochem. Mol. Biol. 2012 |
| WP_007751324.1 | putative 7α-HSDH | *Bacteroides finegoldii* DSM 17565 | Kisiela J. Steroid Biochem. Mol. Biol. 2012 |

| | | | |
|---|---|---|---|
| WP_011860631.1 | putative 7α-HSDH | *Clostridioides difficile* 630 | Kisiela J. Steroid Biochem. Mol. Biol. 2012 |
| NP_810824.1 | putative 7α-HSDH | *Bacteroides thetaiotaomicron* VPI-5482 | Ferrandi Appl. Microbiol. Biotechnol. 2012; Kisiela J. Steroid Biochem. Mol. Biol. 2012; Sherrod Biochim. Biophys. Acta 1977 |
| WP_006236005.1 | 7β-HSDH | *Collinsella aerofaciens* ATCC 25986 | Liu Appl. Microbiol. Biotechnol. 2011 |
| WP_004843516.1 | 7β-HSDH | *Ruminococcus gnavus* ATCC 29149 | Lee J. Lipid Res. 2013 |
| AET80684.1 | 7β-HSDH | *Clostridium absonum*/*Clostridium sardiniense* | Ferrandi Appl. Microbiol. Biotechnol. 2011 |
| EDS07887.1 | 20α-HSDH | *Clostridium scindens* ATCC35704 | Ridlon J. Lipid Res. 2013 |
| WP_107631222.1 | putative 20α-HSDH | *Intestinibacillus* sp. Marseille-P4005 | Current study |
| WP_145772380.1 | putative 20α-HSDH | *Denitratisoma oestradiolicum* | Current study |
| WP_003810233.1 | 20β-HSDH | *Bifidobacterium adolescentis* L2-32 | Doden J. Biol. Chem. 2019 |
| WP_051643274.1 | 20β-HSDH | *Butyricicoccus desmolans* ATCC 43058 | Devendran J. Lipid Res. 2017 |
| WP_051196374.1 | putative 20β-HSDH | *Clostridium cadaveris* AGR2141 | Devendran J. Lipid Res. 2017 |
| WP_027640050.1 | putative 20β-HSDH | *Clostridium cadaveris* AGR2141 | Devendran J. Lipid Res. 2017 |
| BAQ31198.1 | putative 20β-HSDH | *Bifidobacterium scardovii* DSM 13734 | Devendran J. Lipid Res. 2017 |
| WP_024111275.1 | putative 20β-HSDH | *Propionimicrobium* sp. BV2F7 | Devendran J. Lipid Res. 2017 |
| AMB99905.1 | putative 20β-HSDH | *Aerococcus urinaehominis* | Devendran J. Lipid Res. 2017 |
| WP_040693668.1 | putative 20β-HSDH | *Propionimicrobium lymphophilum* ACS-093-V-SCH5 | Ly J. Steroid Biochem. Mol. Biol. 2020 |
| WP_073996553.1 | putative 20β-HSDH | *Arcanobacterium urinimassiliense* | Ly J. Steroid Biochem. Mol. Biol. 2020 |
| WP_025022266.1 | aldo/keto reductase | *Lactobacillus hayakitensis* | Mythen Appl. Environ. Microbiol. 2018 |
| KRM19149.1 | oxidoreductase | *Lactobacillus hayakitensis* DSM 18933 | Mythen Appl. Environ. Microbiol. 2018 |
| KRM65245.1 | oxidoreductase | *Lactobacillus agilis* DSM 20509 | Mythen Appl. Environ. Microbiol. 2018 |
| WP_050611824.1 | aldo/keto reductase | *Lactobacillus agilis* | Mythen Appl. Environ. Microbiol. 2018 |
| WP_019206370.1 | aldo/keto reductase | *Lactobacillus ingluviei* | Mythen Appl. Environ. Microbiol. 2018 |
| KRL88436.1 | organophosphate reductase | *Lactobacillus ingluviei* DSM 15946 | Mythen Appl. Environ. Microbiol. 2018 |
| ASN60792.1 | aldo/keto reductase | *Lactobacillus curvatus* | Mythen Appl. Environ. Microbiol. 2018 |
| WP_085844664.1 | aldo/keto reductase | *Lactobacillus curvatus* | Mythen Appl. Environ. Microbiol. 2018 |
| WP_054778020.1 | aldo/keto reductase | *Lactobacillus saniviri* | Mythen Appl. Environ. Microbiol. 2018 |
| BAN77682.1 | conserved hypothetical protein | *Adlercreutzia equolifaciens* DSM 19450 | Mythen Appl. Environ. Microbiol. 2018 |
| WP_041715350.1 | SDR family oxidoreductase | *Adlercreutzia equolifaciens* | Mythen Appl. Environ. Microbiol. 2018 |
| CDD77593.1 | putative uncharacterized protein | *Cryptobacterium* sp. CAG:338 | Mythen Appl. Environ. Microbiol. 2018 |
| CDD68244.1 | putative uncharacterized protein | *Eggerthella* sp. CAG:368 | Mythen Appl. Environ. Microbiol. 2018 |
| CDB33831.1 | putative uncharacterized protein | *Eggerthella* sp. CAG:209 | Mythen Appl. Environ. Microbiol. 2018 |
| CCY06514.1 | putative uncharacterized protein | *Eggerthella* sp. CAG:1427 | Mythen Appl. Environ. Microbiol. 2018 |
| CDD56334.1 | putative uncharacterized protein | *Bacteroides pectinophilus* CAG:437 | Mythen Appl. Environ. Microbiol. 2018 |
| WP_044925038.1 | short-chain dehydrogenase | *Anaerostipes hadrus* | Mythen Appl. Environ. Microbiol. 2018 |
| WP_008116638.1 | NAD(P)-dependent oxidoreductase | *Bacteroides pectinophilus* | Mythen Appl. Environ. Microbiol. 2018 |
| WP_077326212.1 | short-chain dehydrogenase | *Anaerostipes hadrus* | Mythen Appl. Environ. Microbiol. 2018 |
| | short-chain dehydrogenase | *Clostridiales bacterium* Nov_37_41 | Mythen Appl. Environ. Microbiol. 2018 |
| WP_055161728.1 | short-chain dehydrogenase | *Anaerostipes hadrus* | Mythen Appl. Environ. Microbiol. 2018 |

| | | | |
|---|---|---|---|
| WP_055258939.1 | short-chain dehydrogenase | *Anaerostipes hadrus* | Mythen Appl. Environ. Microbiol. 2018 |
| CCX88977.1 | dehydrogenases with different specificities | *Clostridium sp.* CAG:590 | Mythen Appl. Environ. Microbiol. 2018 |
| WP_009265642.1 | short-chain dehydrogenase | Lachnospiraceae bacterium 5_1_63FAA | Mythen Appl. Environ. Microbiol. 2018 |
| AAB41916.1 | 3α-HSD | *Homo sapiens* | Khanna J. Biol. Chem. 1995 |
| NP_612556.1 | 3α-HSD | *Rattus norvegicus* | Cheng Mol. Endocrinol. 1991 |
| NP_001315544.1 | 3β-HSD/$\Delta^5 \rightarrow \Delta^4$/isomerase type 1 | *Homo sapiens* | Thomas J. Steroid Biochem. 1989 |
| NP_001007720.3 | 3β-HSD/$\Delta^5 \rightarrow \Delta^4$/isomerase type 1 | *Rattus norvegicus* | de Launoit J. Biol. Chem. 1992 |
| NP_001159592.1 | 3β-HSD/$\Delta^5 \rightarrow \Delta^4$/isomerase type 2 | *Homo sapiens* | Rheaume Mol. Endocrinol. 1991 |
| Q04828.1 | AKR1C1 (20α-HSDH) | *Homo sapiens* | Zhang J. Mol. Endocrinol. 2000 |
| NP_001748.1 | Carbonyl reductase 1 (20β-HSDH) | *Homo sapiens* | Morgan Sci. Rep. 2017 |
| AAA30980.1 | 20β-HSDH | *Sus scrofa* | Tanaka J. Biol. Chem. 1992 |
| NP_861420.1 | 11β-HSD type 1 | *Homo sapiens* | Tannin J. Biol. Chem. 1991 |
| NP_058776.2 | 11β-HSD type 1 | *Rattus norvegicus* | Agarwal J. Biol. Chem. 1989 |
| NP_000187.3 | 11β-HSD type 2 | *Homo sapiens* | Wilson J. Clin. Endocrinol. Metab. 1995 |
| NP_000404.2 | 17β-HSD type 1 | *Homo sapiens* | Winqvist Hum. Genet. 1990 |
| NP_002144.1 | 17β-HSD type 2 | *Homo sapiens* | Labrie DNA Cell Biol. 1995 |
| AAC50066.1 | 17β-HSD type 3 | *Homo sapiens* | Geissler Nat. Genet. 1994 |
| AAC15902.1 | 17β-HSD type 10 | *Homo sapiens* | He J. Biol. Chem. 1999 |
